# Dissemination and pathogenesis of toxigenic *Clostridium perfringens* strains linked to neonatal intensive care units and Necrotising Enterocolitis

**DOI:** 10.1101/2021.08.03.454877

**Authors:** Raymond Kiu, Alex Shaw, Kathleen Sim, Harley Bedwell, Emma Cornwell, Derek Pickard, Gusztav Belteki, Jennifer Malsom, Sarah Philips, Gregory R Young, Zoe Schofield, Cristina Alcon-Giner, Janet E Berrington, Christopher Stewart, Gordon Dougan, Paul Clarke, Gillian Douce, J Simon Kroll, Lindsay J Hall

## Abstract

**Background:** *Clostridium perfringens* is an anaerobic toxin-producing bacterium that has long been associated with intestinal diseases, particularly in neonatal humans and animals. More recently, infant gut microbiome studies have suggested an important link between *C. perfringens* and the devastating preterm-associated disease Necrotising Enterocolitis (NEC), but in-depth studies on this pathogen (genomics and mechanistic) are lacking.

**Methods/Materials:** We isolated and whole-genome sequenced 274 infant-associated *C. perfringens* isolates from 5 hospitals across the UK between 2011-2016 (including longitudinal samples from 31 individuals). We performed in-depth genomic analyses, phenotypically characterised pathogenic traits of 10 strains (including 4 *C. perfringens* from NEC patients) and established a novel oral-challenge C57BL/6 mouse infection model for microbe-host studies.

**Results:** Pore-forming toxin encoding genes *pfoA* and *cpb2* were enriched within hypervirulent lineages that exclusively consisted of *C. perfringens*-associated NEC (CPA-NEC) strains, in addition to overabundance of colonisation factors. Importantly, we identified a circulating *C. perfringens* variant, eventually linked to a fatal CPA-NEC case. The variant was detected consistently within 6 individuals in two sister hospitals across a 40-day window, demonstrating for the first time the intra- and inter-hospital dissemination of *C. perfringens*. CPA-NEC isolates were determined phenotypically to be more virulent (linked with overabundance of gene *pfoA*) than isolates obtained from non-NEC preterm babies. In addition, two *pfoA*-positive CPA-NEC *C. perfringens* strains were confirmed to induce clinical inflammatory tissue lesions *in vivo*.

**Conclusions:** Hypervirulent lineages are linked to CPA-NEC, potentially due to the production of pore-forming toxins, coupled with higher metabolic, transmission, and pathogenic capacities. These studies indicate *C. perfringens* is an important bacterial pathogen in preterm infants and highlights the requirement for further investigation into development of intervention and therapeutic strategies.

## Introduction

*Clostridium perfringens* is a Gram positive, anaerobic spore-forming bacterium that is frequently linked to various intestinal diseases in both humans and animals, particularly in neonates. Diseases include gastroenteritis, poultry necrotic enteritis, non-foodborne diarrhoea, and preterm Necrotising Enterocolitis (NEC) ^1^. This gut pathogen is known to secrete >20 toxins, including several pore-forming toxins e.g., β-toxin, perfringolysin O (PFO), NetB and enterotoxin (CPE), which represent primary virulence factors associated with different facets of pathophysiology ^2, 3^. *C. perfringens* also has numerous traits which facilitate intestinal colonisation including rapid proliferation, spore-forming capability ^1,4,5^, and a number of plasmid-encoded virulence factors such as toxins and antimicrobial resistance (AMR) determinants ^6,7^.

*C. perfringens* has low GC content, between 27-28%, and genome sizes often range between 3.0-4.1Mb, with horizontal gene transfer playing a major role in evolutionary adaptation ^7^. Since the first *C. perfringens* strain 13 genome was sequenced and published in 2002, there are currently 275 genomes available on NCBI Genome database (June 2021) ^8^. Although Whole Genome Sequencing (WGS) has advanced our understanding of this gut pathogen, including genomic epidemiology studies for adult-associated gastroenteritis-associated strains, studies of association with other intestinal diseases (and younger age groups) are very limited ^9,10^.

NEC, the most severe and lethal neonatal gastrointestinal (GI) emergency worldwide, is an acquired inflammatory gut disease that manifests as tissue necrosis in the GI tract ^11,12^. NEC typically affects 5-15% very-low-birth-weight (<1,500g) preterm infants, leading to high mortality rates worldwide (~40%), and severe long-term complications including neurodevelopmental delay and short bowel syndrome ^13–16^. Microbial infection represents one of the key risk factors leading to NEC onset ^17,18^. Intestinal pathogens such as *Klebsiella* spp. and *C. perfringens* have been linked to clinical NEC in recent years ^19,20^. Several extensive microbiome-based preterm infant NEC cohort studies have implicated pathogenic bacteria overgrowth prior to NEC, with *C. perfringens* abundance increasing in the gut microbiota several days before NEC diagnosis ^21–23^. Furthermore, *C. perfringens* has been consistently reported since the 1970s as a potential pathogenic agent associated with preterm-NEC (more than any other individual bacterial agent), which has been further supported by *in vivo* studies ^1,24,25^. *C. perfringens*-associated NEC (CPA-NEC) symptoms are often more fulminant and severe compared to ‘classic’ NEC, with infants often requiring surgery, for what has previously been called ‘*C. perfringens* intestinal gas gangrene’^19^.

Here, we describe an extensive genomic study on 274 *C. perfringens* clinical isolates from 5 hospitals across the UK, collected between 2011-2016, some of which were isolated from CPA-NEC patients. We predefined CPA-NEC in this study as definite NEC diagnosis (Bell Stage II/III) with overgrowth of *C. perfringens* prior to NEC onset (Bell’s staging classification system: Bell Stage II/III are considered as definite NEC while Bell Stage I covers non-specific signs) ^22,26^. We probed virulence factors and conjugative plasmids associated with infant/CPA-NEC *C. perfringens* isolates, and traced transmission within and between Neonatal Intensive Care Units (NICUs) via genomic Single Nucleotide Polymorphism (SNP) analysis. We also performed experimental assays on a selection of 10 *C. perfringens* isolates (NEC vs non-NEC) to understand pathogenic strain-level traits. Finally, we developed a *C. perfringens* oral-challenge infection model in C57BL/6 neonatal mice to assay intestinal pathology.

## Results

### Phylogenetic analysis of *C. perfringens* strains reveals distinct preterm-associated lineages

We successfully isolated, cultured and sequenced the genomes of 274 *C. perfringens* isolates from stool samples collected longitudinally from 70 individuals (including 26 isolates from 4 CPA-NEC infants), obtained from 5 UK hospitals between 2011 and 2016 (figure 1A-1C). Of these 274 isolates, 220 were obtained from faecal samples of NICU resident preterm infants while the remaining 54 isolates were cultured from follow-up samples of former NICU infants. There is a ~24% incidence of *C. perfringens* across all infants (table 1). We next reconstructed a core-genome phylogenetic tree of 450 *C. perfringens* strains (including 176 public genome assemblies; figure 1D), which clustered into 6 major lineages (figure 1E). Lineage IV comprised exclusively infant-related isolates (97/97) from hospitals in multiple locations. Other genetically distinct lineages included lineage I (consisting of widely diverse samples and classic human gastroenteritis isolates) and lineage III (including isolates of animal origin 64.3%). Lineage II and V encompassed samples of mixed sources – both humans (101/139; 72.7%) and animals (38/139; 27.3%), with several isolates from poultry Necrotic Enteritis, denoting a potential zoonotic signature. Notably, lineage VI also comprised mostly hospital infant-associated samples (48/59; 81.3%).

**Figure 1.**
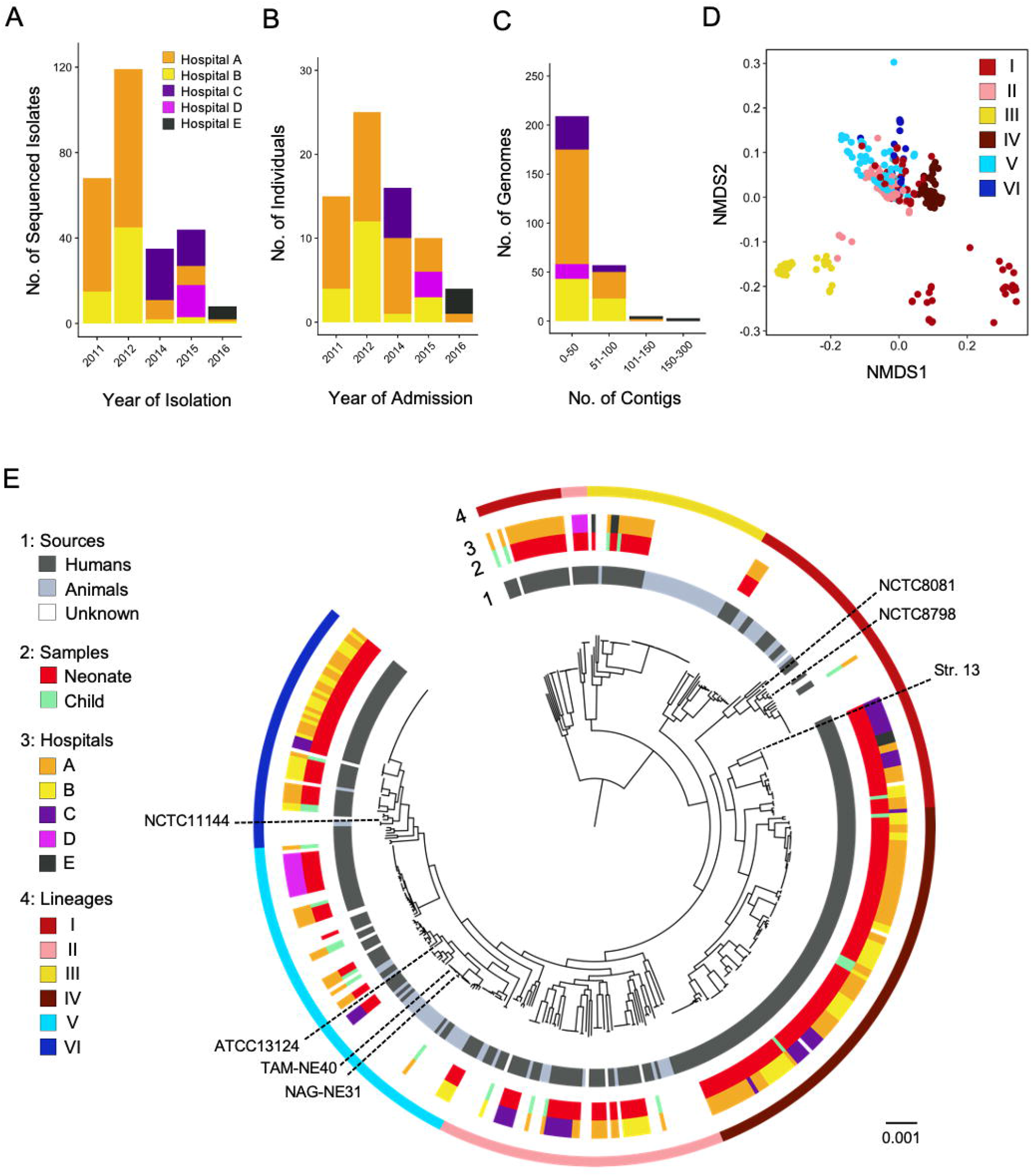
Phylogenomic analysis of 450 *C. perfringens* strains. (A) Year of isolation of the sequenced isolates from 5 hospitals (double-anonymised). (B) Histogram of number of individuals involved in this genomic study in each year (2011-2016). (C) Quality of draft genome assemblies used in this study. Approximately 92% of the genome assemblies used in this study were of high integrity, <100 contigs (short-read draft genome assemblies). (D) Visualising clustering of isolates of various lineages in a Non-Metric Multidimensional Scaling plot based on pangenome binary profiles (gene-presence-absence). (E) Mid-point rooted recombination-free maximum-likelihood tree of 450 *C. perfringens* strains inferred from 23,355 SNPs (from 1,008 single-copy core genes), aligned with metadata including sources, sample hosts, hospitals and lineages assigned using hierBAPS. Black dashed lines, type strain ATCC13124 and several reference genomes were labelled in this tree. Branch lengths are indicative of the estimated nucleotide substitution per site (SNPs).

**Table 1.**
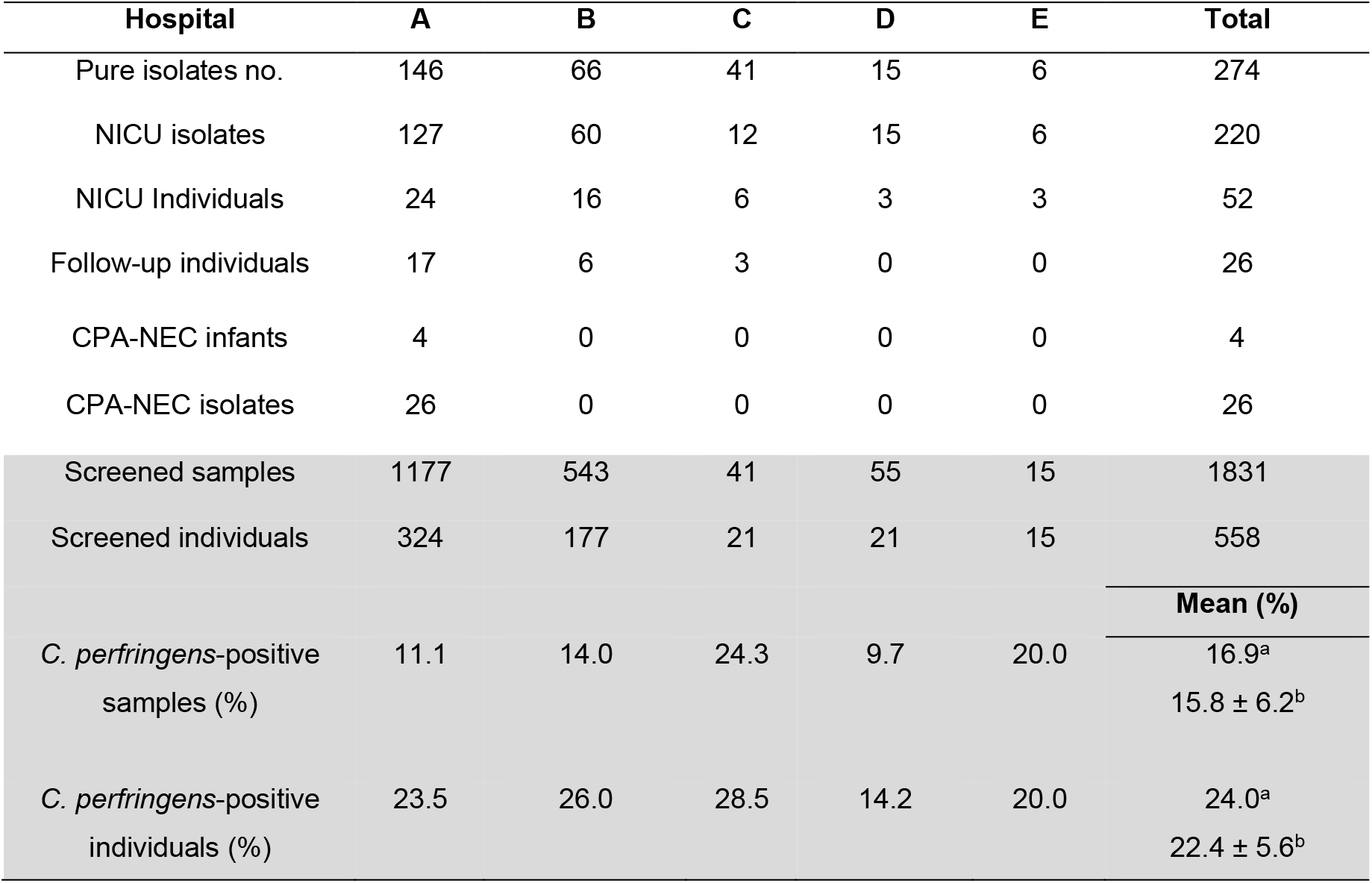
A summary of sample screening, *C. perfringens* isolates and incidence (colonisation data) in this multi-centre collaborative study. Shaded area represents the incidence data. ^a^ Mean (%) calculated from total number of pooled samples across all hospitals. ^b^ Mean ± S.D. (%) calculated from samples across individual hospitals.

We also investigated the pangenome of the 450 *C. perfringens* strains, which revealed a core genome consisting of only 1,008 genes (4.7%), out of a total 21,098 genes (figure S1). On average, ~33% of each individual *C. perfringens* genome (~3.32Mb; ~2,985 genes) are core genes. Furthermore, substantial recombination events were predicted in the core genome of 450 *C. perfringens* strains, which was apparently more common in lineage I, V and VI (figure S2).

### NEC-linked lineages have specific virulence signatures compared to non-NEC lineages

Genome-based virulence profiling was performed on all 274 infant-associated *C. perfringens* isolates (figure 2A). Overall, lineage III isolates carried significantly more virulence-associated genes (median: 17/isolate) than isolates from other lineages, while lineage IV was determined to have less virulence potentials (median: 10/isolate; P<0.05; figure 2B). All CPA-NEC isolates, bar one (25/26), nested in lineages I, II, III, V and VI – which we defined as NEC-linked lineages (P<0.001, significant association with CPA-NEC isolates; figure 1C). The largest single lineage IV (95 isolates from 28 individuals) consisted of ~99% non-CPA-NEC infant isolates – that is to say, a non-NEC lineage (94/95). Comparing the virulence potentials of isolates between NEC-linked and non-NEC lineages, NEC-linked lineages (I, II, III and V) encoded significantly higher virulence gene counts (median: 13/isolate) than non-NEC lineage IV (median: 10/isolate; P<0.0001; figure 1D). Two pore-forming toxin genes *pfoA* (perfringolysin O; PFO) and *cpb2* (β2-toxin; CPB2) were both significantly enriched in isolates of NEC-linked lineages (P<0.0001; figure 1E). Colonisation-associated genes that encode sialidases (*nan*), hyaluronidase (*nag*) and adhesin (*cna*) were also overrepresented in NEC-linked lineages (I, II, III, V and VI) when compared to hypovirulent lineage IV samples (P<0.0001; figure S3).

**Figure 2.**
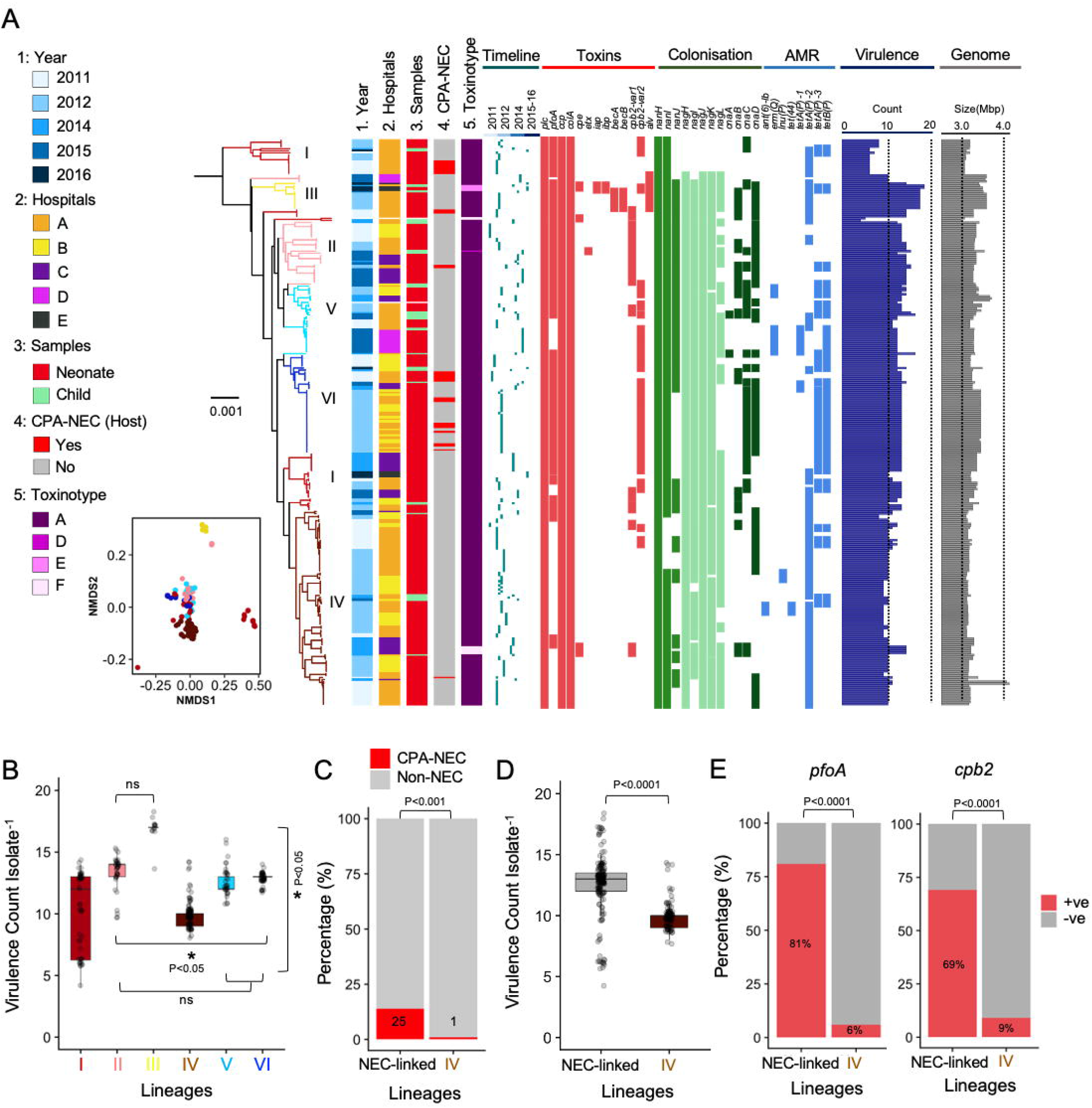
Virulence potentials of infant-associated *C. perfringens* strains. (A) A mid-point rooted maximum-likelihood tree of 276 *C. perfringens* strains inferred from 16,735 core SNPs (predicted recombinant sites removed) aligned with clinical metadata and virulence profiles of each isolate including toxin genes, colonisation-factors and AMR genes, additionally with virulence gene counts and genome sizes indicated. This phylogenetic tree includes ATCC13124 (type strain) and NCTC8679 as reference genomes. Preterm-NEC-associated isolates are labelled in red. Branches are colour-coded according to lineages assigned in Figure 1. The NMDS mini plot represents the clustering of each isolate colour-coded by its lineages. AMR: Antimicrobial Resistance. (B) Virulence count comparison across isolates by lineage. Stats: Kruskal-wallis test, Dunn’s multiple comparison test. Data represented in boxplots. (C) Percentages of isolates from CPA-NEC and non-NEC individuals in NEC-linked lineages (l+ll+lIl+V+VI) vs lineage IV. The numbers in the bars represent the numbers of isolates sampled from CPA-NEC infants. Stats: Fisher’s exact test. Odds Ratio (OR): 15.17. (D) Comparison of virulence count per isolate between NEC-linked lineages (median: 13) and lineage IV (median: 10). Stats: Wilcoxon test. (E) Enrichment analysis of pore-forming toxin genes *pfoA* (OR: 60.80) and *cpb2* (OR: 20.72) between isolates in NEC-linked lineages (l+ll+lll+V+VI; n=179, from 51 individuals) and Lineage IV (n=95, from 28 individuals). Stats: Fisher’s exact test.

### SNP investigation reveals within-host diversity, variant dissemination, vertical transmission, and long-term persistence

In parallel, we investigated circulation of *C. perfringens* isolates via SNP analysis based on longitudinal isolates obtained from individuals as described below. Across all samples from 5 hospitals, SNPs ranged from 0 to ~50 (pairwise SNP comparison; figure 3A). Based on within-host genomic variation of *C. perfringens* strains, we defined 2 SNPs as the SNP threshold to infer transmission (figure 3B). Across 31 infants from three hospitals with ≥2 longitudinal samples (time-points), we identified 9 (out of 31; 29%) individuals who harboured ≥2 *C. perfringens* variants.

**Figure 3.**
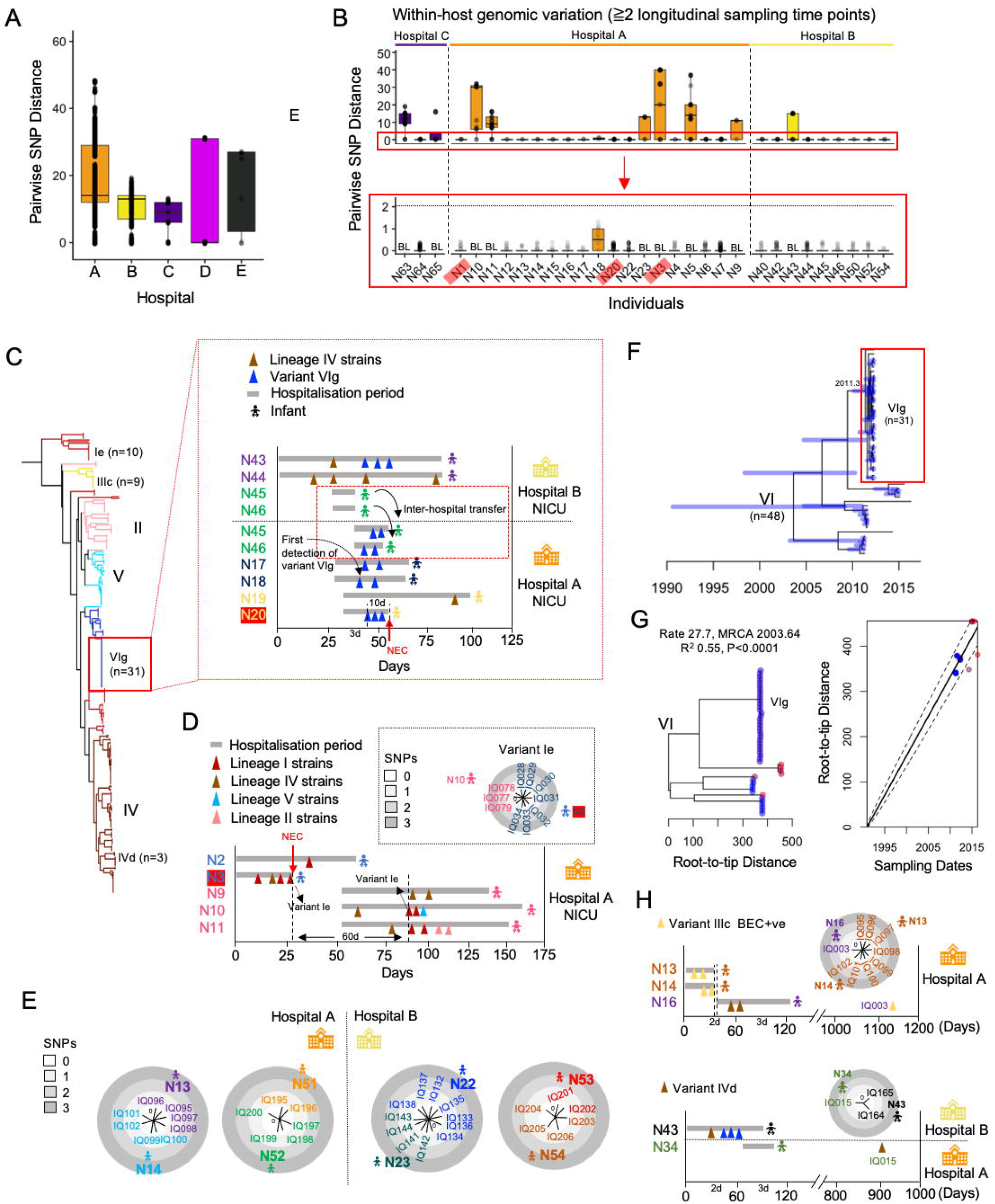
SNP investigation in tracking putative transmission, emergence, and long-term persistence of infant-associated C. *perfringens* strains. (A) Pairwise SNP comparison between *C. perfringens* genomes from 5 hospitals. (B) Within-host SNP variation analysis based on individuals with ≧2 longitudinal time points. Threshold for inferring transmission was defined as ≦2 SNPs. Individuals labelled in red are CPA-NEC patients. (C) Emergence of variant VIg isolates (n=31) that involves 6 individuals in two sister hospitals A and B, based on clinical metadata. Twin pairs are colour-coded. (D) Putative intra-hospital circulation of variant Ie. Variant Ie was not detected in a 60- day window in the infant cohort. (E) SNP distance trees comparison between isolates from twin pairs, indicating potential sources of vertical transmission in the NICUs. (F) Timeline of predicted emergence of variant VIg based on temporal analysis, indicating variant VIg emerged ~12 months before detected in one of the infants. Time bars (year) shown in the bottom. (G) Root-to-tip distance analysis denoting temporal signals of the dataset. (H) Inference of dissemination and long-term persistence of variants IIIc (BEC+ve) and IVd.

Despite apparent within-host-diversity, sub-lineage VIg represented a clonal cluster consisting of 31 isolates (pairwise SNPs ranging from 0-1) obtained from 6 infants residing in both sister hospitals Hospital A and Hospital B NICUs (figure 3C). Clinical metadata linkage indicated that index case of variant VIg (of hypervirulent lineage VI) was detected in infant N18 in Hospital A (figure 3C). Later, this variant was detected in several individuals in Hospital A NICU (N17, N18 and N20), one of whom (N20) was diagnosed with CPA-NEC. Infant N20, who was also diagnosed with a condition pre-disposing to GI mucosal hypoxaemia, subsequently succumbed to NEC 10 days after variant VIg was first detected in the stool. Notably, according to the clinical metadata, each of infants N17 and N18 (index case for variant VIg) had an episode of possible NEC which led to administration of antibiotics (broad-range betalactam + vancomycin) for two days and variant VIg was first detected ~8 days after antibiotic administration in N18. In addition, variant VIg was also detected in individuals residing in Hospital B (across the same 100-day window). This suggests potential spread of the variant between hospitals, whether transferred on/in patients, their parents, or even equipment or healthcare staff. Further support for this hypothesis is indicated by variant Ie, to which NEC patient (N3) who carried variant Ie (index case), representing isolates from single sub-lineage Ie (range: 0 SNPs), succumbed on day 27 (figure 3D); after a gap of 60 days with no detection of variant Ie in any individuals, N10 who was later admitted to the same hospital, also harboured variant Ie suggesting the possibility of horizontal transmission and highlighting potential persistence of *C. perfringens* strains in the NICU environment. Potential vertical transmissions events (from mothers during birth) were also implicated in 4 pairs of twins carrying identical variants (range: 0 SNPs; figure 3E).

We next performed SNP temporal analysis on isolates in lineage VI (n=48) primarily to infer time of emergence of variant VIg and the mutation rate of *C. perfringens* (figure 3F). Emergence of variant VIg was predicted to be ~10 months before the index case N18. Core-genome mutation rate of *C. perfringens* was predicted to be 27.7 SNPs/year/genome, which reflects the diverse genetic nature of this pathogen (figure 3G). We also observed multiple examples of longer-term persistence (figure 3H), where BEC-positive (Binary Enterotoxin of *C. perfringens*) variant IIIc was first detected in infants N13 and N14 (Hospital A), who were both discharged two days before N16 was admitted to the NICU. Notably, we isolated IQ003 as the identical variant IIIc (range: 0 SNPs; sub-lineage IIIc) from N16 stool samples ~1,000 days later (in a follow-on study), suggesting long-term GI persistence. Similarly, we also detected variant IVd (sub-lineage IVd) in preterm N43 at Hospital B, with an identical variant found in preterm N34 ~800 days later, with the infant previously residing at Hospital A, which supports potential environmental transmission and long-term carriage, and/or a hypovirulent variant common in the healthcare environment.

### Virulence-encoding plasmid carriage among infant-related *C. perfringens* isolates

We next investigated plasmid acquisition via *in silico* sequence mapping targeting plasmid-specific conjugative loci, to compare two key *C. perfringens* plasmid families, pCW3 (n=85) and pCP13 (n=111) (figure 4A). Plasmid pCW3 (with *tcp* system) from infant-associated *C. perfringens* isolates encoded toxin genes such as *cpb2, cpe* and iota binary toxin genes *iap* and *ibp*, as well as tetracycline resistance determinants *tetA(P)* and *tetB(P)* (figure 4B). Importantly, adhesin gene *cnaC* was detected in the majority of plasmids (79/85; 92.9%), which may enhance colonisation capacity of these plasmid-carrying strains. NEC-associated isolates from sub-lineage VIg primarily carried pCW3 plasmids in either ~50kb or ~70kb sizes, encoding *tet* determinants (*tetA(P)* and *tetB(P)*), *cnaC*, and toxin gene *cpb2*, with the 50kb plasmids missing ~20 genes due to transposases encoded by *CPR_0630* on both flanks. This plasmid VIg was not detected in strains from other lineages/sub-lineages apart from a lineage V strain IQ074 (infant N10), in which this pCW3 plasmid was determined to be near-identical to those from sub-lineage VIg (range: 2-3 SNPs) (figure 4C). This suggests potential plasmid transmissible events between *C. perfringens* strains, as the NICU stay of preterm N10 overlapped with that of infants carrying sub-lineage VIg strains.

**Figure 4.**
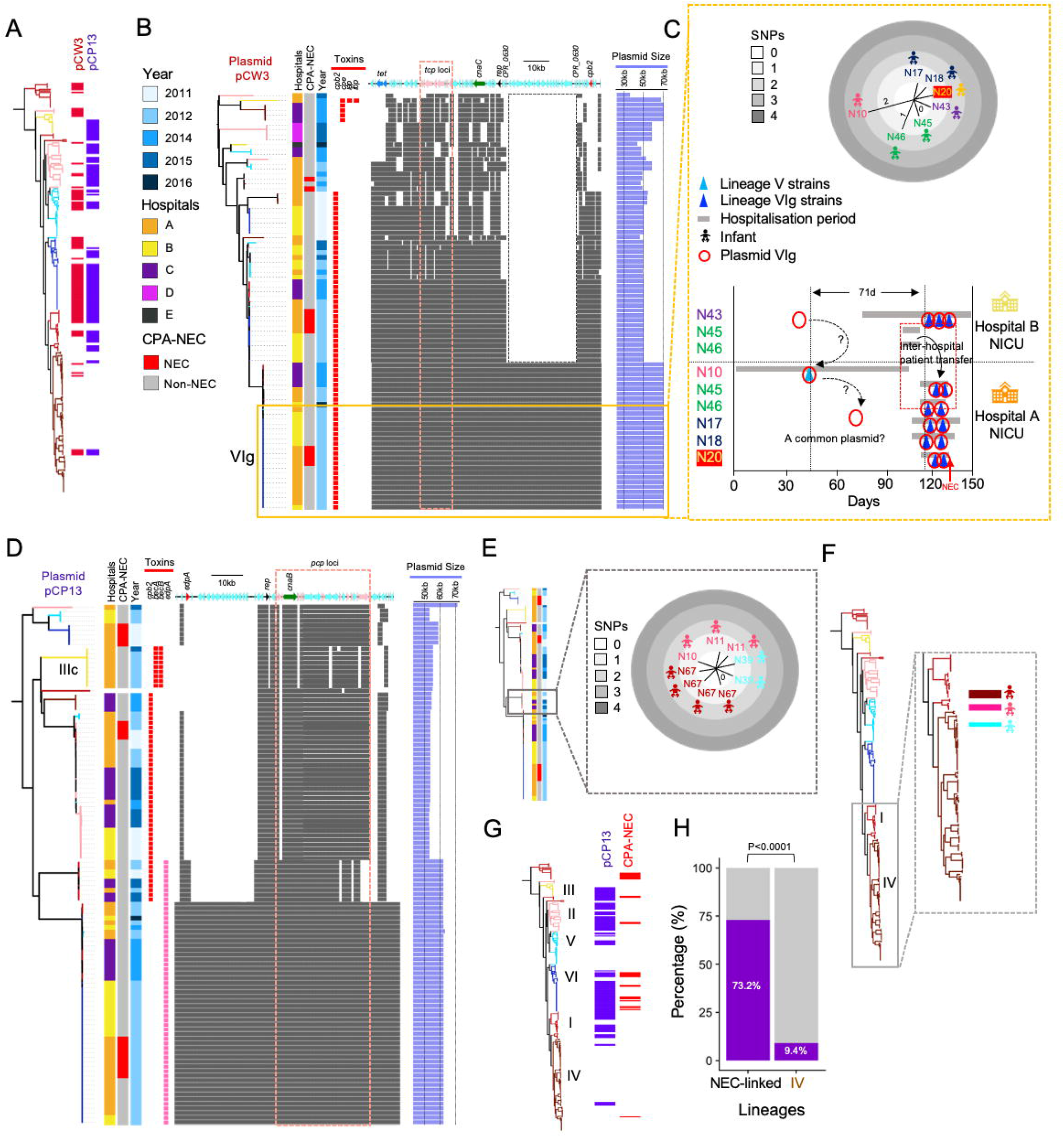
Computational analysis of key conjugative virulence plasmids pCW3 and pCP13 carried by infant- associated *C. perfringens* isolates. (A) Carriage of pCW3 and pCP13 plasmids among infant-associated *C. perfringens* isolates indicated in the phylogeny. (B) Plasmid coding sequences mapped to pLH112 to present a comparative overview of pCW3 plasmids computationally extracted from infant-associated *C. perfringens* isolates (n=85). (C) SNP distance tree of plasmid VIg (top); Probable transmission map of plasmid VIg (bottom). (D) Plasmid coding sequences mapped to pLH112 to present a comparative overview of pCP13 plasmids plasmids computationally extracted from infant-associated *C. perfringens* isolates (n=111). (E) SNP distance tree of near-identical plasmids from 4 individuals (n=9). (F) Host isolates’ position within the phylogenetic tree. These isolates from two major lineages I and IV carried identical plasmids. (G) Carriage of pCP13 plasmids in infant-associated isolates represented in the phylogeny, with CPA-NEC isolates also indicated (NEC). (H) Percentages of isolates carrying ≧1 conjugative virulence plasmid(s), pCW3 and/or pCP13, a comparison between NEC-linked lineages and lineage IV (hypovirulent lineage). Stats: Fisher’s exact test. Odds Ratio: 25.69.

Subsequently, we examined pCP13 plasmid carriage in these infant isolates (figure 4D). The pCP13 plasmids (~45-60kb) in our dataset harboured *cpb2, bec*, and *edpA* (recently identified) toxin genes. Adhesin *cnaB* was detected in all extracted pCP13 plasmids (111/111). Importantly, sub-lineage IIIc strains carried pCP13 plasmids that encoded *becA* and *becB* toxin genes, a binary toxin that has been associated with human gastroenteritis, with further analysis confirming near-identical plasmids from 3 individual infants (figure S4).

Another pCP13 plasmid (~62kb) appeared to be carried by 9 isolates (identical, pairwise SNPs: 0; figure 4E), obtained from 4 individuals, that clustered in two separate lineages (lineages I and IV; figure 4F), confirming the common transferable nature of conjugative plasmids among genetically distinct *C. perfringens* strains. Plasmid pCP13 was frequently carried by strains from all NEC-linked lineages (58.7%; 105/179 isolates; figure 4G). Contrastingly, this plasmid was significantly less common in non-NEC lineage IV, detected only in 6.3% (6/95) of isolates. Overall, conjugative virulence plasmid carriage (either pCW3 or pCP13) was significantly more frequently associated with isolates from NEC-linked lineages (73.2%; 131/179 isolates; P<0.0001) versus hypovirulent non-NEC lineage IV (9.4%; 9/95 isolates; figure 4H).

### Strain-level *in vitro* and *in vivo* characterisation identifies NEC-linked hypervirulent traits

From the genomic data, it was apparent that isolates from NEC-linked lineages encoded significantly more virulence factors than isolates from the non-NEC lineage. Subsequently, we selected 10 *C. perfringens* isolates based on host clinical metadata and categorised these isolates into three groups (table 2): NEC Bell Stage II/III (n=4), NEC Bell Stage I (n=3), and non-NEC (n=3). We performed a series of *in vitro* assays (cell toxicity, survival and colonisation), aiming to link genotype to phenotype for hypervirulent NEC strains (figure 5).

**Table 2.** Clinical information and metadata of the selected 10 *C. perfringens* isolates used for experimental assays. Isolates from Bell Stage II/III patients (n=4) were diagnosed with CPA-NEC.

| Isolate | Phylogenetic Lineage | Hospital | Individual | Bell Stages | NEC day |
| --- | --- | --- | --- | --- | --- |
| IQ146 | II | Hospital A | N24 | II/III | 40 |
| IQ036 | I | Hospital A | N3 | II/III | 24 |
| IQ129 | VI | Hospital A | N20 | II/III | 19 |
| IQ024 | VI | Hospital A | N1 | II/III | 19 |
| LH019 | II | Hospital D | N59 | I | 32 |
| IQ153 | II | Hospital B | N40 | I | 16 |
| LH115 | I | Hospital C | N64 | I | 28 |
| LH043 | V | Hospital D | N61 | non-NEC | / |
| IQ147 | IV | Hospital B | N39 | non-NEC | / |
| IQ133 | IV | Hospital A | N22 | non-NEC | / |

**Figure 5.**
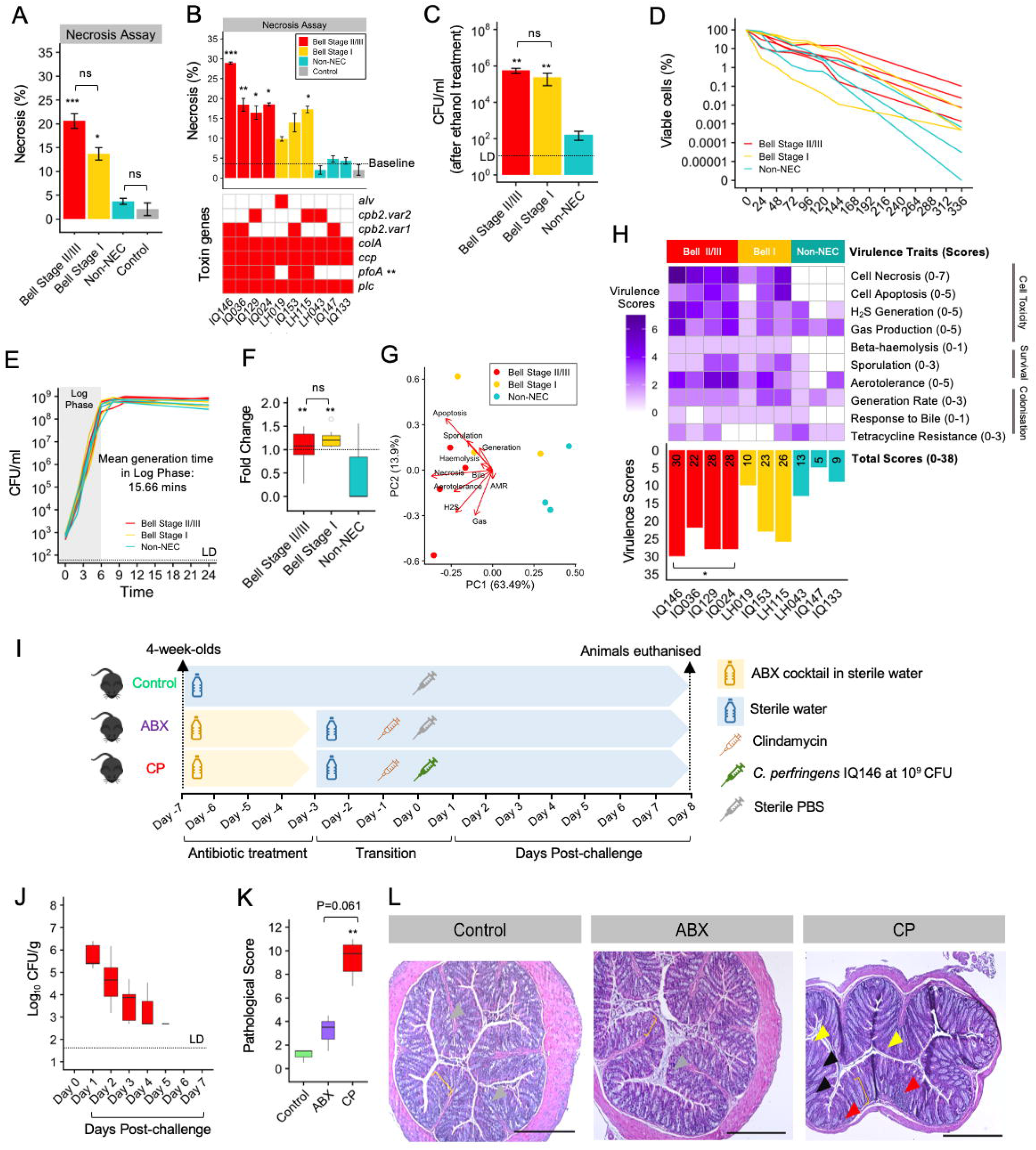
Phenotypic virulence assessment of 10 *C. perfringens* strains. (A) Necrosis percentage of Caco-2 monolayer co-cultured with *C. perfringens* sterile-filtered supernatants after 2h (15% supernatants of in the ratio of 1:2 dilution in the working media). Data: mean±SD (n=3). (B) Necrotic percentage of individual strains aligned with toxin gene profiles. Stats: Point-biserial-correlation test. (C) Sporulation assay. Data: mean±SD (n=3). LD: Limit of Detection. (D) Oxygen tolerance test. Viability of C. *perfringens* strains exposed to ambient air (aerobic) over 14 days. Data: mean (n=3). (E) Growth kinetics. Data: mean (n=3). LD: Limit of Detection. (F) Germination assay. Data: mean±SD (n=3). (G) PCA on overall phenotypic traits. (H) Overall virulence scores comparison. (Data: **** P<0.0001, *** P<0.001, ** P<0.01, * P<0.05. ns: not-significant. Stats: Kruskal-Wallis Test, Dunn’s Multiple Comparison Test for significance.) (I) Scheme of infection model experimental design. (J) Intestinal colonisation of *C. perfringens* monitored over 7 days post-challenge. (K) Pathological scores of murine colons comparison between groups. (L) Representative H&E-stained murine colonic sections of control, ABX and CP groups respectively, showing epithelial changes. Grey arrowhead: normal goblet cells; yellow bracket: the normal colonic crypt length; yellow arrowhead: erosion; black arrowhead: loss of goblet cells; red arrowhead: immune cell infiltration. Magnification: x40, scale bar 50 μm.

To explore *C. perfringens*-associated cellular toxicity, we performed cell necrosis, apoptosis, beta-haemolysis, hydrogen sulphide, generation times, and gas production assays (figure 5). Strains’ supernatants in both Bell Stage II/III (CPA-NEC) and Bell Stage I groups had significantly higher necrotic capacity against Caco-2 epithelial monolayers, when compared to control and non-NEC strains (figure 5A and S5A). This phenotype correlated with presence of the pore-forming toxin gene *pfoA* (P<0.01), suggesting the necrotic potential was primarily due to the effect of pore-forming toxin PFO (figure 5B). Similar findings were also observed with rat intestinal cell line IEC-6 monolayers (figure S5B-S5D). All *pfoA*-encoding isolates (7/7; 100%) also exhibited beta-haemolysis on blood agar plates, confirming the secretion of PFO, whereas non-*pfoA*-carrying strains displayed gamma-haemolysis (haemolysis-free; figure S5E). Bell Stage II/III group strains were also found to produce significantly more gas (figure S5F-S5G) which correlated with hydrogen sulphide (H2S) production (figure S5H-S5I), a known cytotoxic gas against gut epithelial cells (figure S5J).

We next investigated the survivability of *C. perfringens* strains via sporulation (figure 5C and S6A) and aerotolerance assays (figure 5D). Both Bell Stage II/III and Bell Stage I strains were stronger spore-formers than non-NEC strains, while NEC Bell Stage II/III strains were also significantly more aerotolerant at 336 h (after 2 weeks left in ambient air) compared to Bell Stage I and non-NEC strains (figure S6B).

Doubling rates, responses to bile salts and antibiotic resistance capacity were analysed to understand potential colonisation effects. Mean doubling time of all tested strains was 15.66 mins, with no significant differences detected between groups (figure 5E and S7A). Strains from Bell Stage groups had significantly enhanced responses to bile salts, highlighting higher germination capacities in these strains compared to non-NEC strains (figure 5F and S7B). Minimum Inhibition Concentration tests against gentamicin and penicillin (figure S7C) indicated all but two strains were resistant to gentamicin (above MIC breakpoint assigned for *Clostridium* spp.), with all strains susceptible to penicillin. Resistance against tetracycline correlated with the *tet* resistance determinants; with joint presence of both *tetA(P)* and *tetB(P)* correlating significantly with tetracycline MICs (figure S7D).

Principal Component Analysis, based upon phenotypic traits (subjective virulence scoring), revealed that non-NEC strains exhibited a distinctly less virulent phenotypic profile, when compared to NEC-associated strains (figure 5G). The subjective virulence scoring (for ranking purpose and selection of strains for *in vivo* studies) also indicated that Bell Stage II/III strains were significantly more virulent than other groups (P<0.05); IQ146 ranked first (30/38), IQ129 and IQ024 were joint-second (28/38), while IQ147 (5/38) and IQ133 (9/38), both from hypovirulent lineage IV, were the least virulent strains (figure 5H).

Due to the apparent lack of an appropriate *in vivo* intestinal *C. perfringens* infection model, we developed a new oral-challenge murine model. We utilised a 4-day antibiotic cocktail pretreatment prior to one dose of clindamycin 24 hours before oral-challenge of 10^9^ CFU *C. perfringens* IQ146 vegetative cells (strain ranked first in virulence scoring; figure 5I). Colonisation of *C. perfringens* IQ146 was monitored daily post-challenge, which indicated the strain was cleared completely by day 5 (figure 5J). Murine gut microbiome analysis supported the transient presence of *C. perfringens*, with elevated *Clostridium* genus observed in the first three days but absent in both control and antibiotic groups (figure S8A). As expected, antibiotic pre-treatment (ABX) elicited a profound effect on microbiome profiles when compared to the Control group (figure S8B). Significant weight loss was observed on day 2 in the *C. perfringens* group (CP vs ABX and Control; figure S8C), and histology-based pathological scoring indicated the infected group also had mild to moderate colitis (range: 7-11/14) (P<0.01; figures 5K-5L and S8D), when compared to antibiotic and control groups (figure S8E). We observed similar pathological changes in colonic tissues in a modified oral-challenge model, replacing PBS with tetracycline; attempting to prolong the colonisation period using tetracycline-resistant strains IQ146 (Group A) and IQ129 (Group B; figure S9A). Both *C. perfringens* infected groups exhibited higher mean pathological scores compared to the control group (figure S9B). CFU counts indicated that strain IQ129 was able to persist throughout the experimental period, maintaining high 10^7^ CFU/g, while IQ146 was cleared completely before day 5 (in agreement with our previous study; figure S9C). Colonic sections of both infected groups demonstrated similar pathological features, including marked goblet cell loss and immune cell infiltration (figure S9D).

## Discussion

Genomic analysis of preterm infant-associated *C. perfringens* isolates suggests the possible circulation of ‘hyper’-virulent NEC-linked strains and associated virulence-carrying mobile elements within NICUs. Selected NEC-associated isolates were phenotypically determined to be more virulent compared to non-NEC *C. perfringens*, suggesting these strains are capable of initiating pathogenesis in preterm infants.

We assigned 450 *C. perfringens* genomes into 6 major lineages in our phylogenetic analysis, one more than described in a previous smaller study (173 isolates) ^27^. Most lineages, including lineages I, II, III and V, comprised a mixture of diverse genomes (from both human and animal origins), as indicated by extended length of phylogenetic branches (figure 1). In contrast, exclusively infant-associated *C. perfringens* genomes clustered in the monophyletic lineage IV. *C. perfringens* has long been recognised as a potential microbial agent associated with preterm NEC since the 1970s, with two recent metagenomic studies supporting the involvement of *C. perfringens* in microbial-NEC cases ^21,22^.

Currently, public genome databases are biased towards diseased-associated isolates, with only a limited number of *C. perfringens* genomes obtained from healthy individuals (~5%) ^7,27^. As *C. perfringens* can also be considered a commensal bacterium ^28–31^, we sought to understand if there was a genomic ‘signature’ associated with CPA-NEC variants compared to strains found in non-NEC preterm infants. We observed a ‘hypovirulent’ or potentially ‘commensal’ monophyletic lineage (IV), with strains encoding significantly less virulence factors and mobile genetic elements (plasmids), and thus distinct from more pathogenic *C. perfringens*, including CPA-NEC strains (as in lineages I, II, III, V and VI) ^32^. Although these isolates may potentially be hospital-acquired (from our epidemiology analysis), our data suggest distinct *C. perfringens* variants are members of the normal microbiota, and that they may not encode the virulence traits required to cause overt disease such as NEC.

One such genomic signature that appeared enriched in virulent *C. perfringens* isolates, was presence of the gene *pfoA* that transcribes the pore-forming haemolytic toxin PFO. Notably, we observed a direct correlation between the presence of *pfoA* in CPA-NEC isolates (n=4) and significantly enhanced necrotic capacity in intestinal cell lines, when compared to non-NEC/commensal (*pfoA*-negative) isolates. PFO (or, theta-toxin), a typical cholesterol-dependent cytolysin, is known as a pore-forming toxin ^33^, which has been heavily associated with the pathogenesis of myonecrosis (gas gangrene; reported in both animals and humans) and haemorrhagic enteritis in calves ^34–36^. PFO can act synergistically with alpha-toxin in the pathology of gas gangrene, which is a disease that shares a high degree of symptom similarity with CPA-NEC in preterm infants, which has also been called ‘intestinal gas gangrene’^19,37,38^. The mechanistic role of PFO in disease development is currently not well defined (apart from its apparent haemolytic nature), although previous studies have indicated additional modulation of host responses ^34,39–41^. Indeed, our *in vivo* studies suggested a high degree of intestinal immune cell infiltration (and associated pathology) during *C. perfringens* infection. Notably, although previous studies have indicated that *pfoA* is universally encoded in *C. perfringens*, we did not observe this, as demonstrated by lineage IV non-NEC isolates ^42,34^. This prior assumption may be due to skewing in public database for disease-associated strains. Our study therefore highlights the important role of this toxin in disease involvement including CPA-NEC, and it may distinguish between more pathogenic vs. commensal *C. perfringens*.

A concern with more virulent *C. perfringens* strains is nosocomial transmission in at-risk populations, which is a huge issue in related *Clostridioides difficile* ^44,45^. Within hypervirulent lineage VI we observed a circulating variant VIg, which was detected in 6 individuals during their NICU stay, and apparently transmitted between two sister hospitals that regularly exchange patients. This variant also carried the *pfoA* gene and was linked to a fatal case of CPA-NEC in patient N20. Transmission of *C. perfringens* has also been demonstrated in similar confined settings such as within elderly care homes (more vulnerable communities), where non-foodborne outbreaks of toxigenic *C. perfringens* take place frequently ^10,46^. Notably, the index case of variant VIg was detected in an individual who had been treated with broad-spectrum antibiotics, suggesting that creation of a more ‘favourable’ niche may have allowed *C. perfringens* to flourish in the microbiota-depleted gut. The VIg isolate IQ129 was also shown to be a prolific spore-former, a trait which would also provide protection against antibiotics, and evidently possessed colonisation resistance mechanisms, as demonstrated by its detection 8 days after antibiotic treatment ^47^. While 6 infants harboured variant VIg, only N20 developed CPA-NEC and ultimately succumbed to the infection. This preterm had been previously diagnosed with congenital heart disease causing reduced blood flow and oxygen supply to the gut, potentially altering the gut microbiota and barrier permeability ^48–50^. Thus, we postulate that a combination of co-morbidity and microbiota disturbances may lead to overgrowth of certain hypervirulent *C. perfringens* variants and NEC development. Further studies exploring wider microbiome dynamics longitudinal in preterm cohorts, complemented by *in vitro* studies (e.g., model colon systems) are required to tease apart potential mechanisms.

Variant persistence was also inferred longer-term – over two years, indicating that successful initial colonisation within the preterm infant gut may be linked to certain colonisation factors such as *cna*, which were over-represented in more virulent *C. perfringens* strains. Vertical transmission of *C. perfringens* was also implicated in 4 twin pairs, although the assumption would be direct transmission from mother to infant during natural childbirth, but two pairs were born via caesarean section. However, the frequent contact between mothers (parents) and infants is also a likely source of transmission, and we cannot discount the possibility of the hospital environment, transfer of equipment or healthcare staff as additional sources of *C. perfringens*^51,52^. Swabbing in NICUs and regular testing of staff may allow these questions to be answered, alongside profiling stool samples from mothers before and post birth.

The widespread dissemination and extensive persistence of certain strains may partly be attributed to several traits. Our phenotypic studies indicated that hypervirulent *C. perfringens* strains are more oxygen tolerant and have enhanced spore forming and germination abilities, which may facilitate spread between infants and NICUs. This may include standard disinfectant practices (such as 70% ethanol), thus enabling this pathogen to persist in even adverse environments (e.g., sterile settings like hospital), and then germinate once spores enter the infant gut ^53,54^. This ability of certain pathogenic strains (including NEC-associated strains) to persist poses a significant challenge in hospitals and for infection control measures. Our data reinforce the importance of the rigorous hygiene measures are needed in NICUs, especially those wards where extremely low birth weight preterm infants reside who have a heightened risk of developing NEC ^55^. The presence of circulating strains also indicates that routine genomic surveillance of *C. perfringens* may be helpful in NICUs to monitor and prevent circulation of virulent variants.

Certain plasmids (e.g., pCW3 and pCP13) have long been recognised as key mobile genetic elements for transfer of virulence genes, including toxins and AMR determinants in *C. perfringens*^56,57^. We observed 9 *C. perfringens* isolates from both lineages I and IV carrying identical pCP13 conjugative plasmids, suggesting conjugative plasmid transfer to divergent *C. perfringens* strains ^56^. Alongside virulence factors, certain colonisation factors are encoded on these plasmids such as adhesin genes *cnaB* and *cnaC*^57^, with collagen adhesin critical in pathogenesis by conferring additional cell-binding ability in host strains ^58^. Notably, adhesin gene *cnaC*, also required for conjugative transfer, has previously been correlated with virulent poultry-NE *C. perfringens* strains, while a pCP13 plasmid that encodes *cnaB* and *becAB* was recently linked to acute gastroenteritis outbreaks ^57,59–61^. Carriage of these virulence plasmids was found to be more common in hypervirulent isolates from NEC-linked lineages, when compared to commensal isolates, which suggests plasmid encoded virulence traits represent key genomic signatures linked to pathophysiology. The potential plasmid circulation between strains within different wards and hospitals links with recent findings that phylogenetically distinct type-F *C. perfringens* isolates were found to harbour identical CPE- encoding plasmids in a single gastroenteritis outbreak ^10^. However, as our results are based on short-read sequence data (albeit it at very high coverage), further studies using long-read based sequencing and also plasmid extraction and characterisation are required to probe these findings in more detail.

Alongside our in-depth genomic studies, we also explored key virulence traits in ten selected *C. perfringens* strains that are expected to link to intestinal pathology. NEC is associated with fulminant deterioration in preterm infants (often <8h), which may correlate with the rapid proliferation abilities of this bacterium (average *in vitro* doubling time: 15.68 ± 2.33 mins), with subsequent release of toxigenic factors ^4,11,62,63^. Indeed, NEC-linked *C. perfringens* strains had a higher capacity to produce gas, linking to a distinctive NEC symptom – pneumatosis intestinalis (formation of gas cysts in the gut wall), and direct intestinal damage ^11^, with a clinical breath hydrogen test previously used as a NEC diagnostic tool ^64–66^. Furthermore, CPA-NEC strains generated higher concentrations of hydrogen sulphide (H2S), a cytotoxic gas that is associated with other intestinal inflammatory diseases including ulcerative colitis ^67^, and which directly damages the intestinal mucosa, which we also demonstrated *in vitro*^68,69^. This rapid growth and production of necrotic factors in many cases will be prevented by colonisation resistance mechanisms via the resident gut microbiota. However, the preterm gut is known to be significantly disrupted in comparison to full term infants (in which NEC is extremely rare), which may allow overgrowth of *C. perfringens*^1^. The use of probiotics has been proposed as a prophylactic strategy to beneficially modulate the preterm microbiota and reduce incidence of NEC, and previous studies have indicated that supplementation with the early life microbiota genus *Bifidobacterium* correlated with a reduction in *Clostridium* species and associated NEC ^70,71^.

We observed these efficient colonisation resistance mechanisms in our initial animal studies, as adult mice harbouring a complex gut microbiota were refractory to *C. perfringens* infection (data not shown). Thus, we added an antibiotic pre-treatment (5-antibiotic cocktail + clindamycin) regimen in juvenile mice to deplete the resident microbiota, and more closely mimic the clinical scenario in preterm neonates, who are routinely exposed to prophylactic antibiotics in NICUs ^70^. In line with our genomic and *in vitro* studies, we also observed strain-level variations in this model. Strain IQ129, notably a variant VIg, was maintained at high CFU throughout the study period, when compared to strain IQ146, which may link to high spore forming properties (figure S9) enhancing repeated seeding and colonisation in the mouse intestine, and potentially explaining its success as a circulating variant in the NICU. Although we observed different colonisation dynamics between these two strains, similar disease symptoms were apparent including weight loss on day 2 (but no other overt disease symptoms), and mild to moderate intestinal pathology, mimicking some of the pathologies associated with NEC. The pathological changes observed in colonic sections correlated with frequent diarrhoea, which is similar to *Citrobacter rodentium* infection, a model for human enteropathogenic *E. coli*^72,73^. It also appears that immune-mediated pathology plays a role during *C. perfringens* infection, as evidenced by limited mucosal erosions, significant goblet cell reduction and immune cell infiltration (and may link to production of PFO). This implicates the importance of the potential immune-driven aspects of *C. perfringens* intestinal infections, which warrants further investigation. Although CPA-NEC strains were used in this infection model, we did not observe very severe NEC symptoms including complete intestinal necrosis and abdominal distention. This is likely due to differences between mice and humans, and factors such as diet. The standard chow diet may not favour this protein- hungry pathogen, and the use of weaned juvenile mice (rather than very neonatal mice) may lead to more efficient *C. perfringens* clearance (and/or modulation of virulence responses) through immune-mediated pathways, and murine rather than preterm gut microbiota may have impacted infection kinetics. Further optimisation studies, including the use of ‘humanised’ models and/or preterm infant intestinal derived organoid co-culture may allow development of a more clinically relevant NEC-like model ^74,75^.

There are certain limitations associated with this study. Firstly, only 4 CPA-NEC patients, who provided a total of 26 *C. perfringens* isolates, from two sister hospitals, were included. Furthermore, biases may also have been introduced due to the sampling regimen, which in some cases targeted certain individuals across multiple time-points. Within the wider dataset (which included 70 preterms across 5 NICUs) this equates to ~6% NEC incidence. A rate of between 5-15% NEC is often reported, but its incidence varies widely between NICUs, with NEC cases often reported in ‘outbreaks’ that may link to emergence of a specific virulent strain in the hospital environment, a hypothesis supported by our epidemiology analysis ^71,76^. This is particularly important, as previous terminology for the disease - ‘intestinal gas gangrene’ or ‘gas gangrene of the colon’ signals the more severe and fulminant condition of CPA-NEC, as compared to classic NEC, often leading to surgery within 24 hours, and associated with a high mortality rate of up to 78%^19,77^. Larger (and longer) surveillance studies that incorporate numerous NICUs across a wider geographical region would be required to capture a larger number of samples from CPA-NEC diagnosed infants. We also fully recognise NEC as a multifactorial intestinal disease that has been associated with several microbial species including *C. perfringens, Klebsiella* and *Enterococcus* spp ^21,22,78^. Indeed, two preterms had previously been diagnosed with *Klebsiella*-associated NEC in this study, and although they each harboured *C. perfringens* strains, these isolates exhibited a ‘commensal’ genomic signature, were members of the hypovirulent lineage IV, and did not encode *pfoA*. Future work is required to pinpoint the mechanistic role of the virulence factors identified, such as virulence gene *pfoA*, which could be explored using *pfoA*-negative hypovirulent strains tested alongside hypervirulent *pfoA*-positive strains in the oral-challenge murine model.

Overall, we have identified hypervirulent lineages of *C. perfringens* linked to CPA-NEC, with isolates harbouring significantly more virulence factors including pore-forming toxin PFO and plasmids linked to *in vitro* and *in vivo* pathogenic traits. Dissemination and persistence of hypervirulent *C. perfringens* variants was observed between and within preterm infants and hospitals, highlighting the potential value of routine surveillance and enhanced infection control measures. A novel murine infection model can now be further used to study CPA-NEC infection mechanisms and test preventative new measures against *C. perfringens*.

## Methods

### Clinical samples and bacterial isolation work

We conducted a retrospective genomic analysis on *C. perfringens* isolates obtained from faecal samples of 70 neonatal patients admitted to NICUs at Hospital A, Hospital B, Hospital C, Hospital D and Hospital E respectively in the UK between February 2011 and March 2016 (Table S1). Dates of hospital admission and transfers were extracted electronically. Faecal samples collected were stored at −80°C freezer prior to experimental process to isolate *C. perfringens* using ethanol-shock method (50% ethanol in Robertson’s Cooked Meat Media), followed by plating on Fastidious Anaerobic Agar supplemented with defibrinated sheep blood and 0.1% sodium taurocholate; alternatively, faecal samples were plated directly on Tryptose-Sulfite-Cycloserine Egg Yolk Agar (TSC-EYA) prior to 37°C anaerobic incubation for 18-24 h ^79^. Multiple or single distinct colonies on the plates were purified and maintained as pure isolates in autoclaved Brain Heart Infusion (BHI) broth with 30% glycerol for cryopreservation at −80°C.

### Microbiology, whole-genome sequencing and *de novo* genome assemblies

Pure isolates were cultured anaerobically overnight at 37°C in BHI broth (~10-15 h) for genomic DNA extraction using either phenol-chloroform extraction method as described previously, or FastDNA SPIN Kit for soil according to manufacturer’s instructions (Hospital E samples only; MP Biomedicals) ^80^. WGS of each isolate sample was performed on Illumina HiSeq 2500 to generate 101/125 bp paired-end reads as described previously ^81^. Isolate samples from Hospital E were sequenced on Illumina NextSeq 500 to generate 150 bp paired-end reads.

Raw sequence reads (FASTQ) were quality-filtered with fastp v0.20.0 prior to *de novo* genome assembly using SPAdes v3.14.1 at default parameters ^82^. Contigs of <500 bp were filtered in each genome assembly before subsequent analyses. Genome assembly statistics were generated via sequence-stats v0.1 (Table S2) ^83^.

### Phylogenetic analyses

A total of 450 *C. perfringens* genome assemblies, including 176 public genomes retrieved from NCBI Genome database (remaining 274 are novel draft genomes generated in this study), passed contamination checks including Average Nucleotide Identity (ANI) via fastANI v1.3 ^84^ (>95% ANI vs type strain ATCC13124), GC content (in between 27-28%) and 16S rRNA sequences via BACTspeciesID v1.2 ^85^ (>99% nucleotide identity vs type strain ATCC13124) and were used for further analyses. A core-gene alignment (1,008 single-copy core genes) of 450 *C. perfringens* genomes based on Prokka v1.13 annotation of coding sequences (GFF) was generated using Roary v3.12.0, at blastp 95 % identity, adding option -s (do not split paralogs), and options -e and -n to generate core gene alignment using MAFFT v7.305b ^86–88^. Maximum-likelihood phylogenetic trees were constructed based on the core-gene alignment (678,155 bp) via RAxML-NG v0.9.0 ^89^ using general time-reversible GAMMA (GTR+G) model with 100 bootstrap replicates, prior to filtering recombinant sites using ClonalFrameML v1.12 at default parameters ^90^. Infant-associated phylogenetic tree was reconstructed in similar fashion, with two reference genomes ATCC13124 and NCTC8679 in addition to the 274 novel genomes. Lineages and sub-lineages were assigned using R library RhierBAPS v1.0.1 sequence-clustering algorithm based on recombinant-free core-gene alignment ^91^. Non-metric Multi-dimensional Scaling (NMDS) clustering was performed using R package *vegan* based on gene-presence-absence matrix of the pangenome ^92^. Temporal analysis was performed with R package *bactdating* v1.0 using ClonalFrameML outputs, mixedgamma model, with 1e^6^ MCMC iterations (nbIts) and added argument ‘useRec = T’ for main function *bactdate* ^93^. The effective sample size (MCMC) of the inferred parameters α, μ and σ were computed to be >180. Node labels (dates) were extracted from dated tree using FigTree v1.4.4. Tree annotation was performed via iTOL v4.0 ^94^.

### Toxinotype assignment and virulence profiling

Toxinotype A-G was assigned to each *C. perfringens* sample via TOXIper v1.1 ^95^. Virulence- related genes and colonisation factor sequence search was performed via ABRicate v1.0.1 with options --minid=90 and --mincov=90 based on in-house sequence databases (Table S3). ResFinder v4.0 database was used via ABRicate for profiling antimicrobial resistance genes ^96^.

### Single Nucleotide Polymorphism (SNP) analysis

SNPs were extracted from core-gene alignment (recombinant sites removed) with snp-sites v2.3.3 and snp-dists v0.7 was utilised for computing pairwise SNP distances ^97,98^. Transmission analysis was limited to hospital-associated strains (from 70 individuals) and within closest genetic distance identified by phylogenetic topologies and pairwise SNP distances within non-recombinant core-gene alignment of 1,008 single-copy core genes (nested in the same sub-lineage and ≦2 SNPs genetic distance were criteria to infer transmission). Probable transmission dynamics were predicted based on clinical metadata of the patients and the time of the individuals’ staying within the hospitals. Persistence was initially predicted on the basis of isolates of different individuals nested within the same sub-lineages; SNP-distance was further explored to determine the exact genetic distance to infer transmission.

### Plasmid analysis

Plasmid analysis was limited to hospital-associated *C. perfringens* genomes. Conjugative plasmids, both pCW3 and pCP13 families (known to encode multiple virulence factors), were extracted based on the identification of the *tcp* (pCW3 plasmids; n=12) and *pcp* (pCP13 plasmids; n=20) genes in the conjugative systems as described previously via ABRicate v1.0.1 ^56,57^. Briefly, *tcp* and *pcp* loci, and plasmid replication protein gene *rep* were comprehensively searched on 276 genomes; only single contigs within genome assemblies comprising >5 conjugative genes (*rep* is compulsory) were extracted and assumed as functional conjugative plasmids for further analyses. Predicted genes were mapped to reference plasmids LH112 (the largest plasmid size in both families) to allow comparison of plasmid contents across both pCW3 and pCP13 plasmid families. Further virulence factors were identified via ABRicate v1.0.1 with in-house databases as described in previous section. Plasmid sequences were aligned with MAFFT v7.305b, SNP distance was compared via snp-dists ^88^. Easyfig v2.2.2 was utilised for visualisation of plasmid sequence comparison ^99^.

### Cell line maintenance

For IEC-6, Frozen stocks were resuscitated/resuspended in pre-warmed (37°C) media Dulbecco’s Modified Eagle Medium (DMEM, Thermo Fisher Scientific), supplemented with 10% Foetal Bovine Serum (FBS; Thermo Fisher Scientific), 4mM L-glutamine (Thermo Fisher Scientific) and 0.1 U/ml bovine insulin (Sigma-Aldrich). Cells were immediately transferred to T25 sterile culture flasks (25cm2; Corning) and incubated at 37°C in 5% CO2 incubator for 24h. Spent media was removed after 24-48h, replaced with fresh warm media as described and incubated until confluency was reached. For making cell line stocks, cells were grown to confluency (70-80%) before trypsinisation (1% trypsin for 10 min at 37°C) and cell density adjusted to approximately 1×10^6^ cells/ml. 5% of DMSO (cryoprotectant) was added to 1ml aliquots before transferring to −80°C freezer for 24h. After 24h, stocks were stored to − 196°C liquid nitrogen for long-term storage (cell banking). Cell line passages 6 – 20 were used, cells were split at ratios 1:10-20 for each passage. Cells were counted on haemocytometer (Neubauer Chamber). For Caco-2, cells were resuscitated and maintained as described above except the working medium – DMEM supplemented with 20% FBS. Cells were split at ratio 1:5-10. Cell line passages 30-45 were used in experiments.

### *In vitro* phenotypic characterisation

#### Necrosis assay

IEC-6 cells (passage 6-20) were seeded at 20,000 cells/well in 96-well plate (tissue-treated). After 72h 5% CO_2_ incubation, approximately 56,000 cells per well (firm monolayer) were subjected to experiments. Briefly, phenol-red DMEM was removed, cells were washed twice gently and replaced with sterile phenol-red free DMEM supplemented with 1% FBS. Diluted sterile-filtered (0.22μm) bacterial supernatants (in ratio 1:4) were added into each well at 5% total volume followed by 2h 5% CO_2_ incubation. Necrosis measurement was performed using CytoTox-One Homogeneous Membrane Integrity Assay according to manufacturer’s instructions (Promega). For Caco-2 cells (passage 30-45), cells were seeded at 20,000 cells/well. After 72h 5% CO_2_ incubation, a confluent monolayer was formed. Protocol follows as described, with an exception that supernatants were added in 25% of total volume.

#### Apoptosis assay

Cell seeding follows the above description. After 3h 5% CO_2_ incubation, caspase activity was measured using Caspase-Glo 3/7 Assay according to manufacturer’s instructions (Promega). Staurosporine (1μM) was used as positive control.

#### Growth kinetics assay

Bacterial stocks were checked for purity on BHI agar prior to experiments. Single colonies were picked and cultured in BHI broth anaerobically overnight (approximately 14-16h) to reach stationary phase (approximately 1 × 10^9^ CFU). Confluent cultures were diluted 1- million-fold (10^6^) at the start of the 24-h experiment, estimating 10^3^ starting CFU/ml/ well. Growing cultures were homogenised by inverting 5 times before taking 100μl of total liquid for plating. Serial dilutions were performed with sterile PBS in sterile Eppendorfs before drop-plating using Miles and Misra method on BHI agar ^100^. For each time point and sample, 3 technical replicates were drop-plated for statistical accuracy. This experiment was performed in three independent cultures (n=3) for each isolate. Plates were incubated at 37°C in anaerobic cabinet for 15-20h before CFU counting. Plates of highest dilutions with CFU were selected for maximum accuracy.

#### Gas production assay

Autoclaved gas vials filled with 30ml sterile BHI were pre-reduced overnight in anaerobic chamber prior to experiment. Overnight cultures were diluted to approximately 10^3^ CFU/ml each vial and air-sealed with butyl septum and aluminium crimp, then moved to 37°C incubator. Briefly, gas produced by each culture was measured by USB pressure transducer connected to a 3-way stopcock – inserting a 23G hypodermic syringe needle (attached to a 10-ml syringe) into the sealed vial headspace, the gas pressure readouts were indicated on the display. By withdrawing the syringe plunger, the headspace pressure returned to ambient pressure as indicated by a zero reading ^101^. Gas volume was indicated on the syringe barrel after the headspace pressure returns to zero. Each reading was then recorded every hour for an extended period of 10h. Measurements were carried out in a fume hood.

#### Hydrogen Sulphide tissue toxicity assay

Sodium hydrosulfide (NaHS) was used as a sulfide donor as an equivalence of H2S in this study ^102^ Concentration of soluble NaHS was prepared ranged from 0.0001% to 1% in PBS, sterile-filtered (0.22μm) and stored at 4°C prior to experiment. Cell seeding (IEC-6) was performed as described, with all NaHS solution being added to working volume (200μl) at 5% each well. Supernatants were harvest after 4h 5% CO2 incubation. CytoTox-One Homogeneous Membrane Integrity Assay was used to measure potential cell toxicity according to manufacturer’s instructions (Promega).

#### Hydrogen Sulphide assay

Overnight pure cultures were inoculated in 20ml fresh sterile BHI in 50ml tubes at 1% inoculum and followed by 8h anaerobic incubation (negative control *Bifidobacterium longum* was inoculated at 5% and incubated for 20h). H_2_S test strips (also known as lead acetate test strips: H_2_S detection range 5-400ppm) were attached in each tube to visually determine the amount of H_2_S produced by each strain using blackening score (1 to 10). NaHS was added to BHI to serve as positive control as it produces H_2_S when dissolved in liquid (which gives a blackening score of 10).

#### Beta-haemolysis test

BHI agar supplied with 5% sheep blood was used to identify PFO expression of *C. perfringens* isolates. Briefly, *C. perfringens* was streaked on the blood agar and incubated anaerobically for overnight (<20h). As PFO is known produce beta-haemolysis in blood-supplemented media (complete lysis of blood cells), therefore isolates with clear transparent halos formed around the colonies (indicating lysis of blood cells) visualised were defined as PFO-positive isolates.

#### Oxygen tolerance assay

Pure cultures were grown anaerobically to confluency for 24h in BHI. Cultures were spotted in a dilution series onto BHI agar supplemented with 0.1% sodium taurocholate. Spotted plates were dried and incubated under ambient (aerobic) conditions at room temperature (21°C) for specified time periods (24h-336h) before being returned to anaerobic chamber for determination of CFU count. All CFU were counted after 12-15h anaerobic incubation. Cultures (spotted plates) that were not exposed to oxygen acted as controls for each strain. Viable percentage was shown after comparing with control cultures.

#### Sporulation assay

Pure cultures were induced to sporulate using the modified Duncan-Strong Medium (at pH 7.8) for 24h anaerobic incubation unless otherwise indicated ^103^. Sporulated cultures were treated with sterile-filtered 70% ethanol for 4h to eliminate all remaining vegetative cells and spotted on BHI agar supplemented with 0.1% taurocholate (a potent germinant) for spores.

#### Bile salt assay

Ethanol-treated sporulated pure cultures were serially diluted and plated on BHI agar and BHI agar supplemented with 0.1% bile salts (taurocholate, cholate, chenodeoxycholate and deoxycholate), followed by anaerobic incubation for >24h. CFU were enumerated, and fold-change was calculated compared with CFU on BHI plates without bile salt supplementation.

#### Minimum Inhibitory Concentration assay

The MIC of each antibiotic against *C. perfringens* strains was determined based on the modified method of microdilution technique for antimicrobial susceptibility testing ^104^. MIC is defined as the lowest antibiotic concentration that inhibits the visible growth of *C. perfringens* after 24h-incubation.

Sterile-filtered antibiotics were pre-made in stocks at desirable concentration prior to storage at −20°C. Bacterial culturing was described previously. 96-well plate was utilised for this assay – approximately 10^4^ CFUs of each strain was added into each well (sterile BHI) containing a different antibiotic concentration (double dilution) accordingly. BHI without antibiotic served as a positive control for growth and supplemented with thiamphenicol at 15μg/ml served as negative control. Microplate reader (FLUOstar Omega) was used to measure liquid turbidity at 595nm (absorbance) to determine whether there is microbial growth in each well after 24h anaerobic incubation. Absorbance >0.5 is considered as significant growth.

#### Subjective virulence scoring scheme

A subjective/relative virulence scoring system was employed for the sole purpose of ranking/comparing isolates to reflect the exhibited virulence in the *in vitro* assays. Virulence scores were assigned according to the clinical importance to potential disease pathology with cell necrosis the highest (7), followed by cell apoptosis (5), hydrogen sulphide generation (5), gas production (5), aerotolerance (5), sporulation (3), generation rate (3), antimicrobial resistance (3), response to bile salts (1) and beta-haemolysis (1; as it is ‘binary’ comparison). Virulence outcomes were compared semi-quantitatively, albeit subjectively allocating highest scores to most ‘virulent’ strains, less scores for relatively less ‘virulent’ strains (same scores if relatively similar), and 0 for strains with negligible outcomes.

#### *In vivo* studies

##### Ethics and licence

All animal experiments and related protocols described were performed under the Animals (Scientific Procedures) Act 1986 (ASPA) under project licence (PPL: 80/2545) and personal licence (PIL: I7382F677) and approved by Home Office and UEA FMH Research Ethics Committee. Animals are monitored and assessed frequently during studies for physical condition and behavior. Mice determined to have suffered from distress were euthanised via ASPA Schedule 1 protocol (CO_2_ and cervical dislocation). Trained and qualified animal technicians carried out animal husbandry at UEA Disease Modelling Unit (DMU).

##### Animals and housing

C57BL/6 wild-type female mice (juvenile mice: three weeks old), obtained from DMU were used in animal experiments. Animals were bred and housed in DMU barn under specific pathogen-free conditions and moved to DMU infection suite prior to the study. During infection study, mice were housed with autoclaved bedding (and cage), food (stock pellets) and water with 12h light cycle (12h of light and 12h of darkness). Cages were changed in laminar flow cabinet.

In-house wild-type C57BL/6 3-4 weeks old female mice (after weaning) were treated with a five-antibiotic cocktail that comprises kanamycin (0.4mg/ml), gentamicin (0.035mg/ml), colistin (850U/ml), metronidazole (0.215mg/ml) and vancomycin (0.045mg/ml) in autoclaved drinking water for 3-4 days administered ad libitum.

Drinking water was then switched to sterile water, and 48h later mice were orally gavaged with 150mg/kg of clindamycin. After 24h, mice were challenged with 10^9^ CFU of *C. perfringens* in 100μl and all mice were closely monitored for signs of disease symptoms including significant weight loss (>20%) that will require euthanisation. Tetracycline was supplemented (0.001mg/ml) in drinking water post oral challenge in tetracycline model.

##### Bacterial strains and growth

Bacterial stocks were recovered on BHI agar for purity checks each time and subsequently cultured in BHI broth overnight to reach confluency (10^9^ CFU/ml, approximately 14-17h). Bacterial pellets were washed and re-suspended in sterile PBS before feeding the mice via oral gavaging using 20G plastic sterile feeding tube.

##### Faecal sample collection and CFU enumeration

Faecal samples were collected in sterilized tubes daily and stored at −80°C until further analysis. Serially diluted faecal mixtures were plated on fresh TSC agar and CFU enumerated <24h anaerobic incubation. Pitch black colonies were counted as *C. perfringens* colonies.

##### Tissue processing and H&E staining

Intestinal sections were fixed in 10% neutral buffered formalin for <24h and followed by 70% ethanol. Tissues were processed <5 days in 70% ethanol using an automated Leica Tissue Processor ASP-300-S and embedded in paraffin manually. Sectioning was performed using a microtome (5-μm-thick sections) and left overnight for samples to air-dry prior to staining and further analysis. H&E (haematoxylin and eosin) staining was performed subsequently for structural imaging of intestinal samples.

##### Pathological scoring

All colonic section images were examined and graded single-blinded by L.J.H. The histological severity of infectious colitis was graded using a pathological scoring system (0-14; denoting increasing severity) based on three general pathological features: (1) inflammatory infiltration (0-4), (2) epithelial hyperplasia and goblet cell loss (0-5), and (3) mucosal architecture (0-5) as described previously ^105^. The overall score was the sum of each component score. An overall pathological score of 1-4 indicates minimal colitis, 5-8 as mild colitis, 9-11 as moderate colitis, above 12 as marked/severe colitis.

##### Microscopy imaging

Bright-field microscopy was performed using Olympus BX60 with a microscope camera Jenoptik C10 with ProgRes CapturePro software v2.10.

#### Microbiota analysis

Genomic DNA of murine faecal samples were extracted, sequenced and analysed following the protocol described previously ^60^. Briefly, DNA was extracted from samples using FastDNA Spin Kit for Soil (MP Biomedicals) following manufacturer’s instruction, while extending bead-beating step to 3 min ^70^. Next, DNA extracts were subject to 16S rRNA Illumina MiSeq sequencing library preparation, amplifying V1+V2 regions of the 16S rRNA gene prior to paired-end sequencing at 2 × 300bp ^70^. Sequencing raw reads (FASTQ) were firstly merged using PEAR ^106^, then underwent quality and chimera filtering via QIIME v1.9.1, subsequently OTU assignment using SILVA_132 database at 97% similarity (Table S4). BIOM output of OTU tables were read using MEGAN6 ^107,108^.

#### Graphing and statistical methods

Various statistical graphs including pie chart, line graphs, bar plots and box plots were drawn in R v3.6.3, using R libraries *tidyverse, ggplot2* and *ggpubr* ^109,110^. Pangenome heatmap was generated via phandango ^111^. NMDS plots were drawn using R library *vegan* ^92^. Statistical tests (two-sided) were carried out via R base packages *stats* (Fisher’s exact test, Kruskal-Wallis test and correlation test) and *FSA* (Dunn’s test) ^110^.

## Ethical approval

Faecal collection from NNUH and Rosie Hospital (BAMBI study) was approved by the Faculty of Medical and Health Sciences Ethics Committee at the University of East Anglia (UEA), and followed protocols laid out by the UEA Biorepository (License no: 11208). Faecal collection from Imperial Healthcare NICUs (NeoM and NeoM2 studies) was approved by West London Research Ethics Committee (REC) under the REC approval reference number 10/H0711/39. In all cases, doctors and nurses recruited infants after parents gave written consent. Ethical approval for the SERVIS study was approved by the North East and N Tyneside committee 10/H0908/39, and signed parental consent obtained from every parent. We have anonymised identifiers of patients, hospitals and associated clinical data.

## Data access

Sequencing reads for *C. perfringens* isolates generated in this study have been deposited in the European Nucleotide Archive (ENA) (https://www.ebi.ac.uk/ena) under project PRJEB25762. Raw 16S rRNA amplicon sequences are available under project PRJEB28307. In-house databases for sequence-search and scripts for running R libraries are available at: https://github.com/raymondkiu/Infant-Clostridium-perfringens-Paper

## Author Contributions

Conceptualisation, R.K., A.S, K.S, G.D. (Glasgow), J.S.K. and L.J.H.; Methodology, R.K., A.S, K.S, E.C, J.M., C.A., S.P., Z.S. and L.J.H; Software, R.K.; Validation, K.S., A.S, L.J.H.; Formal analysis, R.K.; Investigation, R.K., K. S., A.S. and L.J.H. Resources, D.P., G.B., J.E.B., G.R.Y., C.S., G.D. (Cambridge), P.C.; Data Curation, R.K., A.S, K.S., C.A.; Writing - Original Draft Preparation, R.K. and L.J.H.; Writing - Review and Editing, R.K., A.S., K.S., Z.S., J.E.B., C.S., J.S.K, L.J.H.; Visualisation, R.K.; Supervision, J.S.K. and L.J.H.; Project Administration, R.K.; Funding Acquisition, J.S.K and L.J.H.

## Acknowledgments

This research was supported in part by the NBI Computing infrastructure for Science (CiS) group through the provision of a High-Performance Computing (HPC) Cluster. L.J.H. is supported by Wellcome Trust Investigator Awards 100974/C/13/Z and 220876/Z/20/Z; the Biotechnology and Biological Sciences Research Council (BBSRC), Institute Strategic Programme Gut Microbes and Health BB/R012490/1, and its constituent projects BBS/E/F/000PR10353 and BBS/E/F/000PR10356. We thank the sequencing team at both Wellcome Trust Sanger Institute and Quadram Institute Bioscience for genome sequencing. We would like to express our appreciation to our colleagues, Dr Fred Warren, Prof Simon Carding, and former colleagues Shabhonam Caim, Dr Lukas Harnisch, Dr Benjamin Kirkup at Quadram Institute Bioscience (QIB) for their discussion on the data and assistance in various trainings associated with this work.

