## Supplementary Figures for "Dissemination and pathogenesis of toxigenic *Clostridium perfringens* strains linked to neonatal intensive care units and Necrotising Enterocolitis"

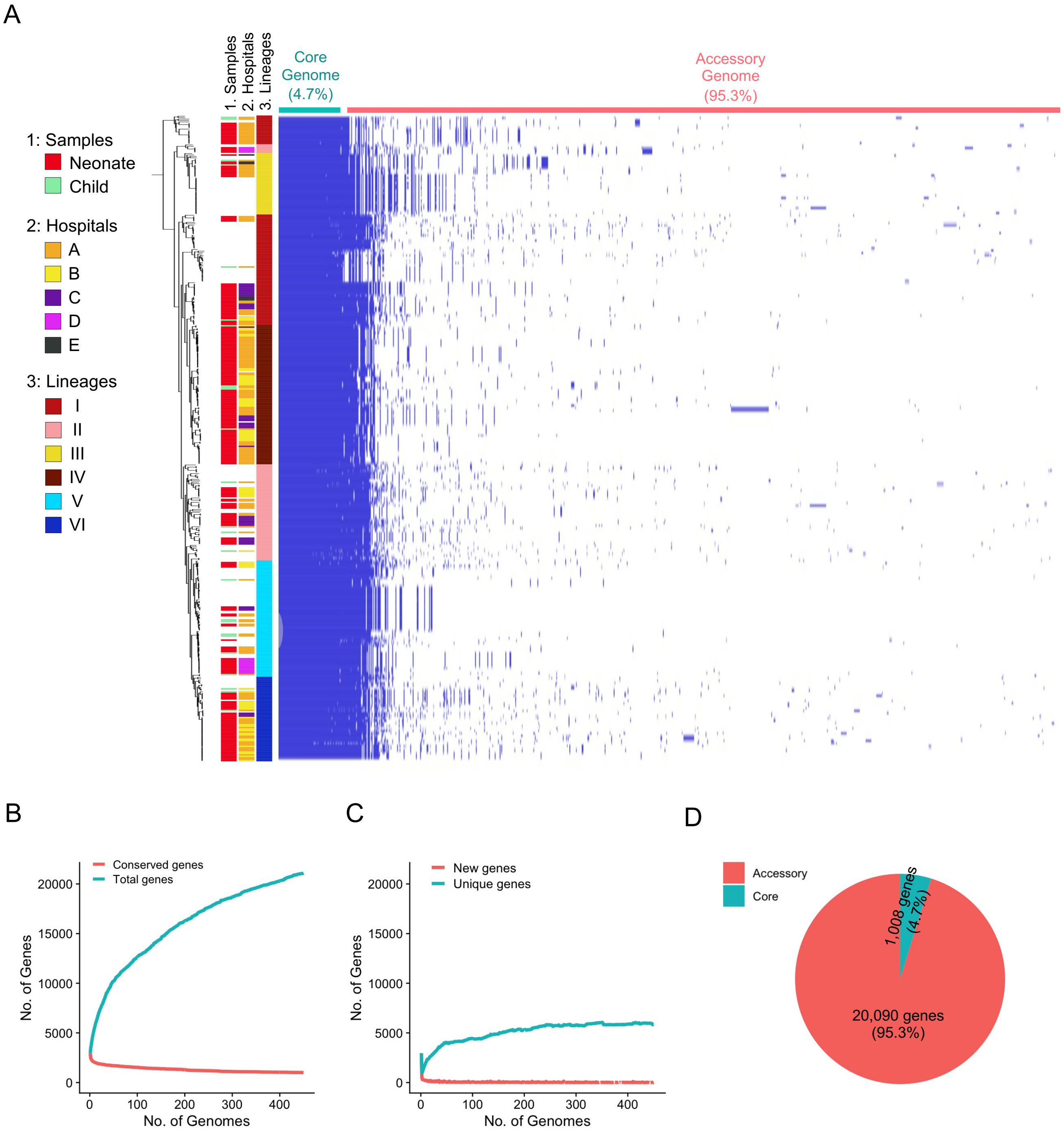


**Figure S1. Pangenome-wide analysis on 450 *C. perfringens* genomes.** (A) Linearised pangenome visualisation of 450 *C. perfringens* genomes in binary heatmap, aligned with clinical metadata and lineages. A total of 21,098 genes were predicted in this pangenome, of which 1,008 genes (4.7%) are core genes (present in 100% of the genomes) while 20,090 genes (95.3%) are accessory (present in at least one genome). (B) Visualisation of gene accumulation in pangenome computation process, conserved genes vs total genes. (C) Visualisation of gene accumulation in pangenome computation process, new genes vs unique genes. Approximately 27 new genes were predicted at the end of pangenome reconstruction, denoting an open nature of this pangenome, towards a tendency of adding more new genes. (D) A pie chart showing the proportion of core genes vs accessory genes in the pangenome.

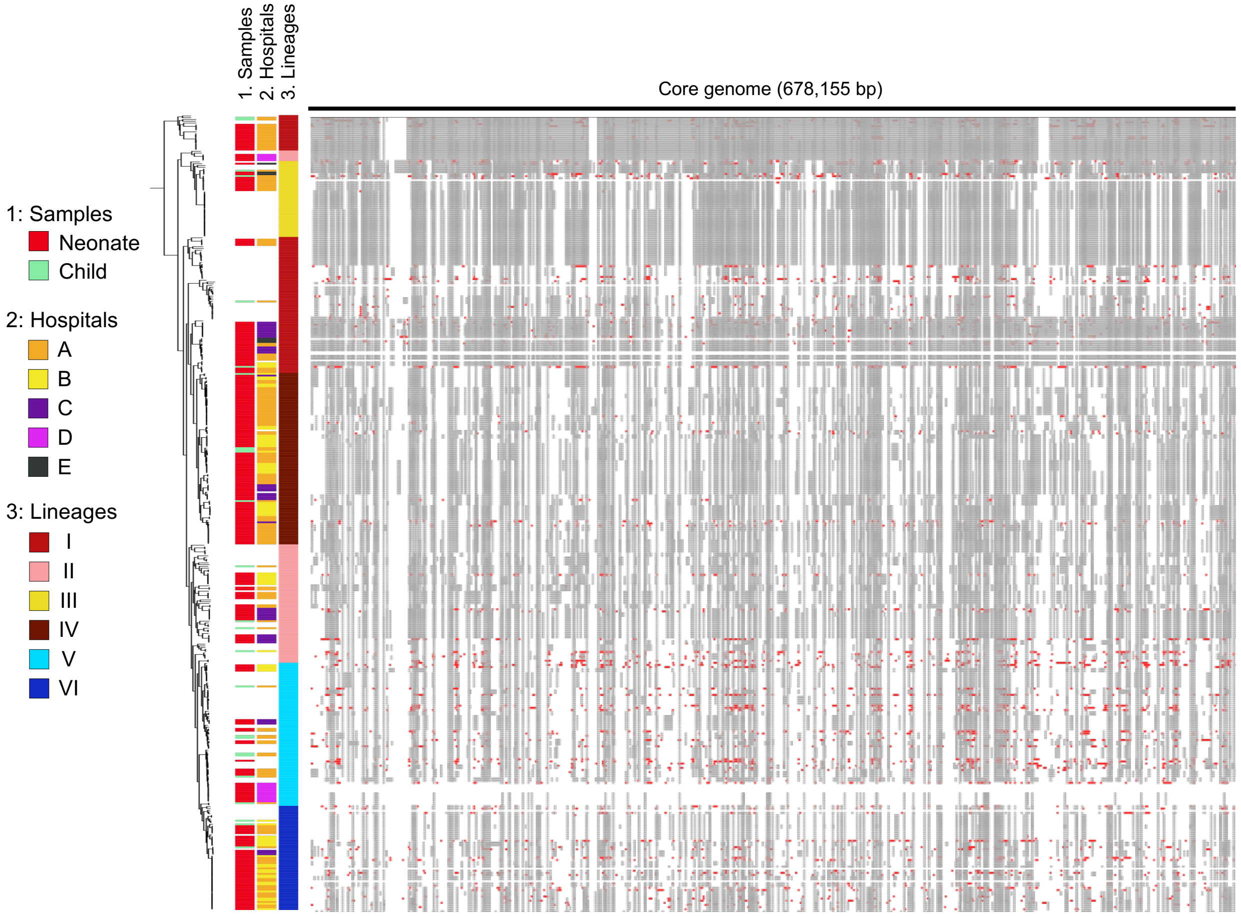


**Figure S2.** Recombination events inferred in the core genome (1,008 genes) of 450 *C. perfringens* strains (678,155 bp). Recombination events in *C. perfringens* core genome were predicted using ClonalFrameML. Ancestral recombinant sites are represented in grey regions while predicted extant recombinant sites in red spots. Apparently, extensive recombination events were observed in lineages I, V and VI.

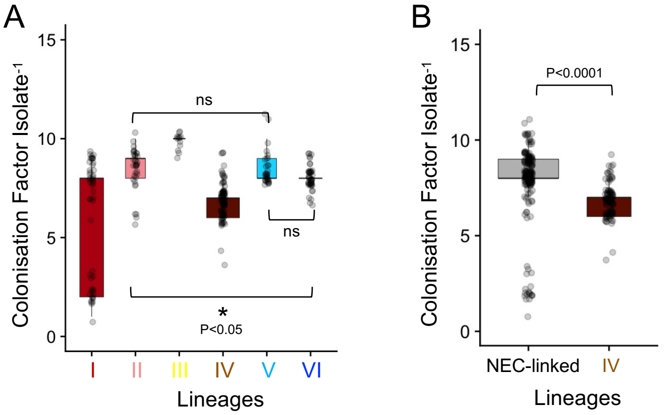


**Figure S3.** (A) Colonisation factor comparison across all isolates by lineage. Stats: Kruskal-wallis test, Dunn's multiple comparison test. (B) Colonisation factor comparison between NEC-linked lineages and lineage IV. Stats: Wilcoxon test. Data represented in boxplots.

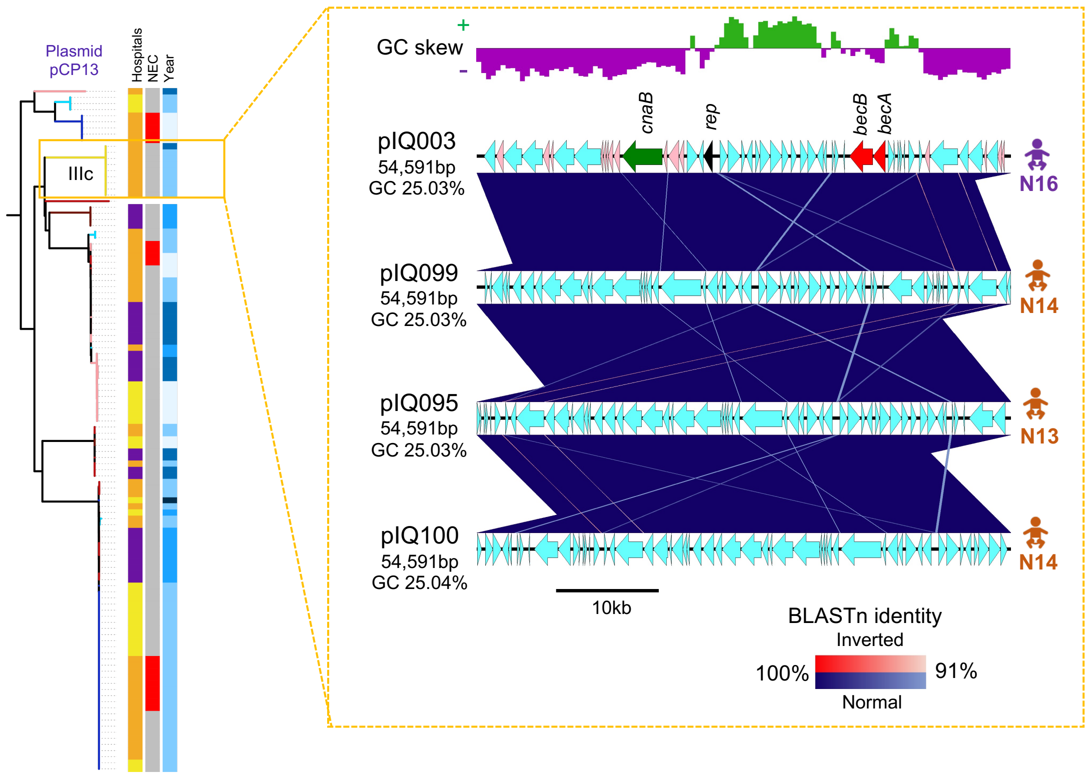


**Figure S4.** Comparative analysis of near-identical plasmids from sub-lineage IIIc isolates.

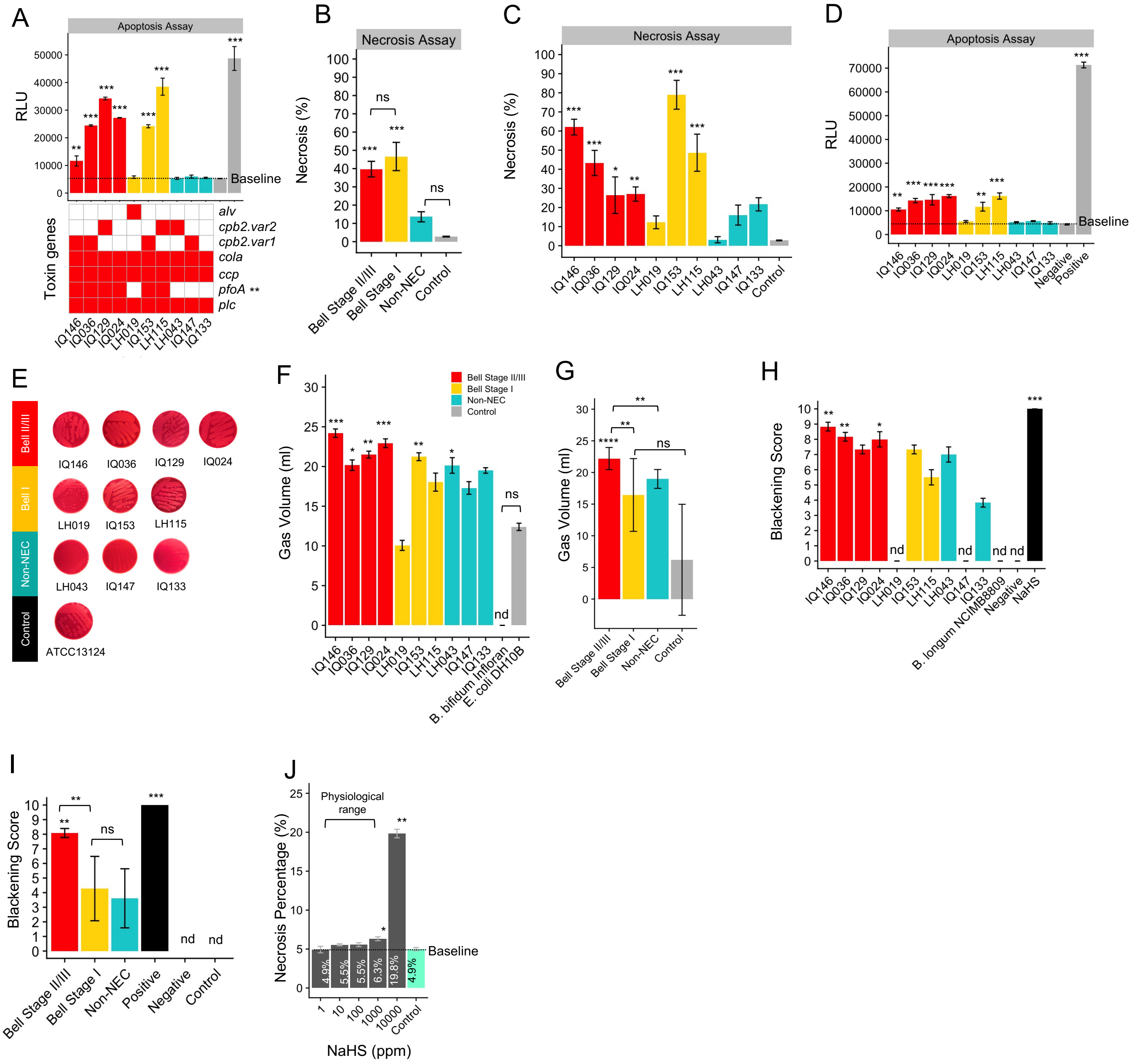


**Figure S5. Cell toxicity assays.** (A) Apoptosis assay on Caco-2 cell lines aligned with toxin gene profiles. Data: mean±SD (n=3). Stats: Point-biserial-correlation test. (B) Necrosis assay on IEC-6 epithelial monolayer in two separate experiments (3 technical replicates each). (C) Necrosis assay on IEC-6 epithelial monolayer, 1:4 dilution in two separate experiments. (D) Apoptosis assay on IEC-6 epithelial monolayer (two separate experiments. RLU: Relative Luminescence Unit. (E) Beta-haemolysis assay. (F) Accumulated gas production of *C. perfringens*. Confluent cultures were inoculated at $10^3$ CFU/ml (exception: *B. bifidum* and *E. coli* at $10^6$ CFU/ml) and sealed in 30ml-sterile air-tight gas vials (50ml) in anaerobic chamber prior to experiment. All cultures and equipment were pre-reduced overnight before experiment. Measurements were taken every hour and the experiments stopped at 10h. nd: Not Detectable. Data: means ± SEM (n=3). (G) Gas production assay, comparison between groups. Data: mean±SD (n=3). (H) Hydrogen sulphide ($H_2S$) visual measurement test. Blackening scoring - 1: slight black stain on the strip; 5: 50%black strip; 10:100% black strip. The degree of blackening was used as a $H_2S$ concentration/production gauge. nd: Not Detectable.(I) Hydrogen sulphide assay, comparison between groups. nd: Not Detectable. Data: mean±SD (n=3). (J) Necrotising effect of NaHS ($H_2S$ donor) on IEC-6 monolayer system at 2h-incubation.

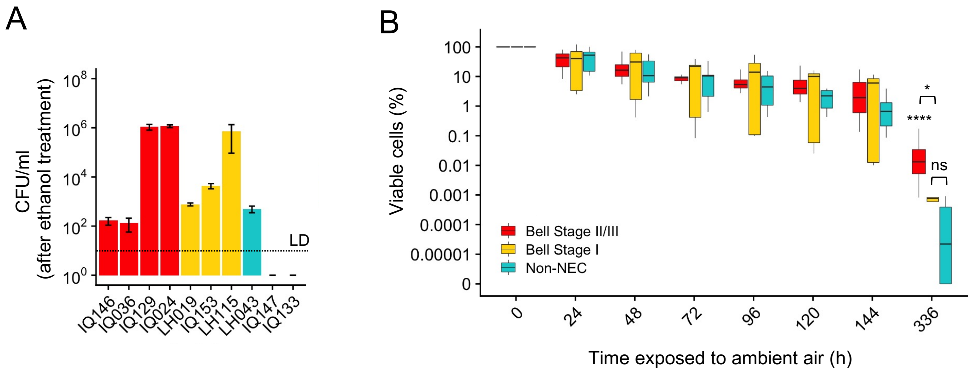


**Figure S6. Survivability assays.** (A) Sporulation capacity test. Vegetable *C. perfringens* strains were sporulated in Duncan-Strong media for 24h prior to 70% ethanol treatment for 4h to eliminate vegetative cells. Spores were then germinated on BHI supplemented with taurocholate, while CFU were counted after 24h anaerobic incubation. Strains IQ147 and IQ133 were repeated with prolonged 48h Duncan-Strong sporulation albeit with negative outcome, no CFU was observed. Data: mean±SD (n=3). (B) Comparisons of *C. perfringens* viability (%) after exposure to ambient air at 8 time-points across experimental period according to clinical groups. For all time-points, no statistically significant differences were found in between diseased/non-diseased groups except time-point 336 h (n=3). (Data: mean±SD; **** P<0.0001, *** P<0.001, ** P<0.01, * P<0.05. LD: Limit of Detection. Stats: Kruskal-Wallis Test, Dunn's Multiple Comparison Test for significance.)

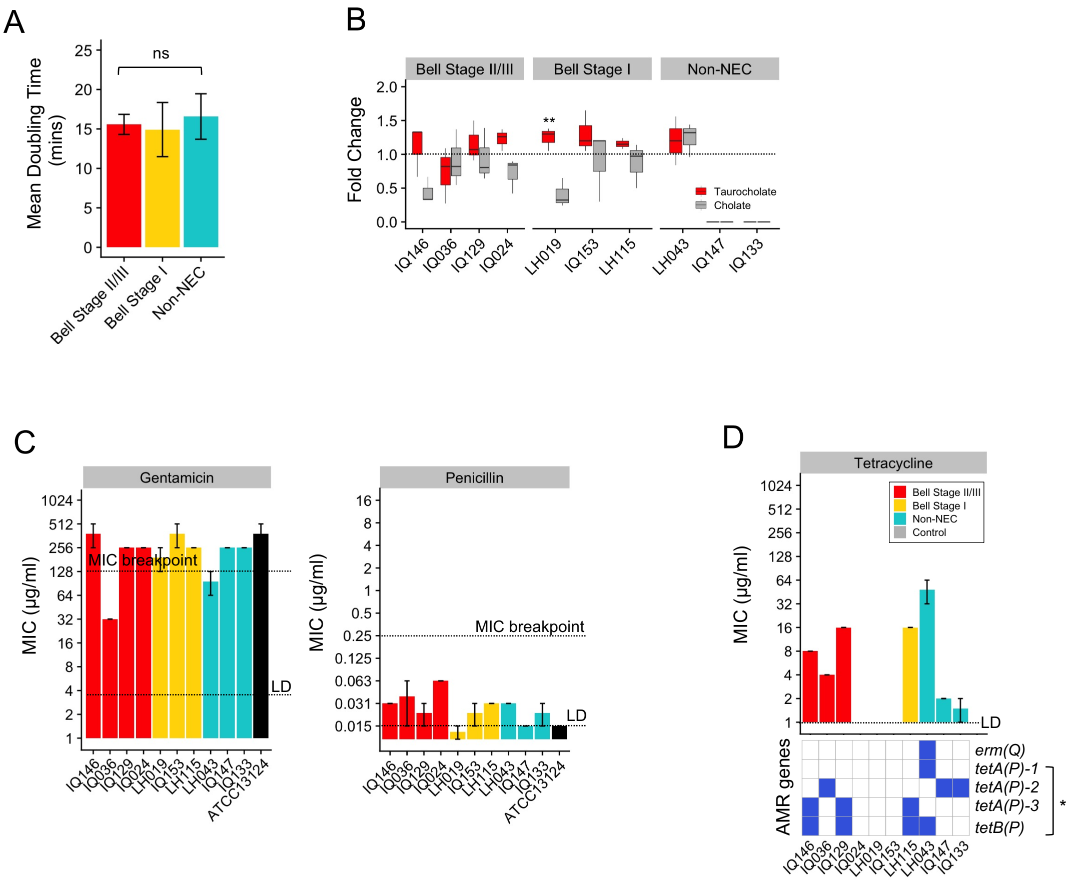


**Figure S7. Colonisation assays.** (A) Mean generation time comparison in clinical groups. ns: not significant. (B) Germination potency (fold-change) comparison. (C) MIC of antibiotics Gentamicin and Penicillin against *C. perfringens*. Type strain ATCC13124 was used as a control strain (two separate experiments). (Data: mean±SD; **** P<0.0001, *** P<0.001, ** P<0.01, * P<0.05. LD: Limit of Detection. Stats: Kruskal-Wallis Test, Dunn's Multiple Comparison Test for significance).

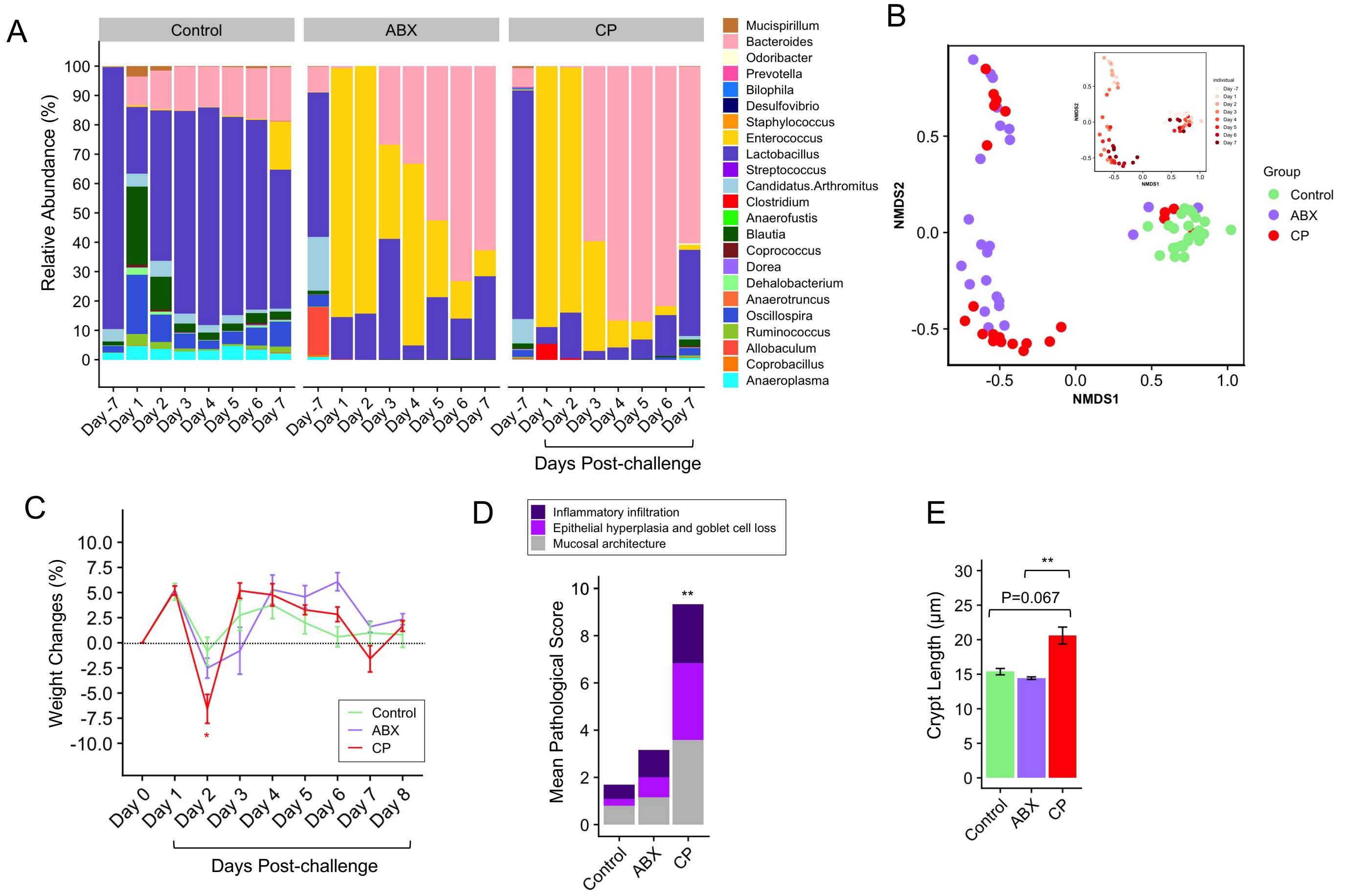


**Figure S8.** Oral-challenge mouse model of *C. perfringens.* (A) Gut microbiome kinetics on genus level (n=3). (B) NMDS plot on gut microbiome kinetics over pre- and post-challenge periods. (C) Daily weight changes of mice over experimental period. (D) Mean pathological scores graded according to three key pathological features (total 0-14): inflammatory infiltration (0-4), epithelial hyperplasia and goblet cell loss (0-5), and mucosal architecture (0-5). (E) Comparison of colonic crypt length of each group. (Data: mean±SD; **** P<0.0001, *** P<0.001, ** P<0.01, * P<0.05. Stats: Kruskal-Wallis Test, Dunn's Multiple Comparison Test for significance).

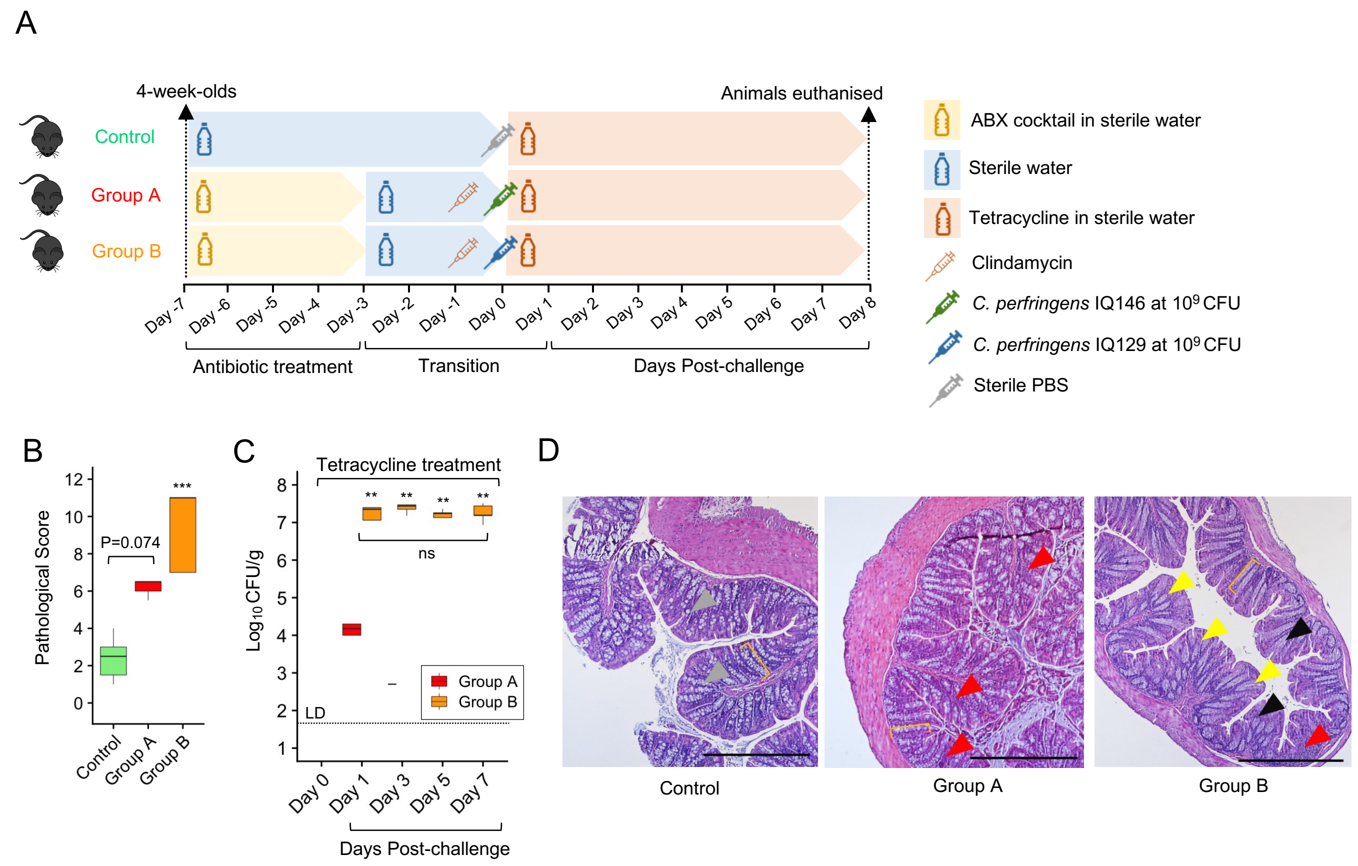


**Figure S9.** Tetracycline oral-challenge mouse model of *C. perfringens.* (A) Scheme of experimental design. (B) Pathological scores of murine colons of each group. (C) Intestinal colonisation of *C. perfringens* strains monitored over 7 days post-challenge. (D) Representative H&E-stained murine colonic sections of Control, Group A and Group B respectively, showing epithelial changes. Grey arrowhead: normal goblet cells; yellow bracket: the normal colonic crypt length; yellow arrowhead: erosion; black arrowhead: loss of goblet cells; red arrowhead: immune cell infiltration. Magnification: x40, scale bar 50 µm. (**** $P<0.0001$, *** $P<0.001$, ** $P<0.01$, * $P<0.05$. Stats: Kruskal-Wallis Test, Dunn's Multiple Comparison Test for significance).
